# Layers in the sand: The genetic imprint of migration, culture, and Indus craft in the Thar desert

**DOI:** 10.1101/2025.09.17.676769

**Authors:** Rishabh Jain, Sachin M. Rathod, Pankaj Jha, Deepika Jangir, Mohammed Faruq, Ganga Nath Jha, Mitali Mukerji

## Abstract

The Thar Desert of northwest India, despite its harsh ecology, has sustained settlement of ancient crafts and pastoral communities. Their persistence provides a unique opportunity to study how migration, ecology, and culture have shaped genetic diversity. We analyzed genome-wide SNP data from 176 individuals across eight occupational communities along with global, Indian populations and diverse ancient genomes. Population history, ancestral migration, population structure, demography, admixture, and founder effects were elucidated using diverse population genetic statistical methods. The Thar groups occupy an intermediate position on the Indian north–south cline. Pastoralists and artisans (woodcarvers and Persian gold embossers) with West Eurasian lineages, while potters and performers align with southern clines. Uniparental data confirmed heterogeneous Indian lineages. Gene-culture was evident from the lactase persistence allele being higher in pastoralists but lower in gold-embossers despite shared ancestry. Noteworthy, the populations retain a high frequency of the SLC24A5 allele, associated with lighter skin pigmentation in Europeans, despite the desert environment. Demographic analyses indicate admixture of 60–80 GBP and strong founder effects in certain groups, particularly tie-and-dye and Persian migrant artisans ∼500–600 years ago. aDNA comparisons confirmed continuity with Indus-periphery and historical South Asian populations. The genetic landscape of the Thar is a palimpsest shaped by successive layers of settlement, migration, and cultural continuity. By establishing the first genomic baseline of Thar’s craft and pastoral communities, this study shows how ecology and endogamy with population history shape distinct genetic landscapes. These findings provide essential context for studying genetic risk, adaptation, and human resilience in extreme environments.

## Introduction

Northwest India lies at a historic crossroad of ancient trade and migration routes connecting South Asia, Central Asia, and the Middle East.^1^ Particularly, the Thar Desert region is characterised by resident populations from diverse ethnic groups with distinct socio-cultural practices, including endogamy, nomadism, and pastoralism. Kin-based occupational continuity is a hallmark of the Thar communities, which are renowned for their vibrant traditional crafts, including pottery, woodcraft, leatherwork, textiles, music, and various dance forms.^2–6^ Many of these practices can be traced back to ancient times, particularly the Indus Valley Civilization (IVC, c. 3300–1300 BCE), which thrived in the adjoining geographical regions encompassing Harappa and Mohenjo-daro in the west, Kalibanga in the north, and Dholavira and Lothal in the south of the Thar Desert.^7–10^ Recently, Harappan remains have been identified in the Thar Desert, indicating that the civilization may have had a broader geographical distribution than previously recognized.

The Thar Desert today is acknowledged as the world’s most densely populated desert. Evident from the continuity of craft, farming, and pastoralism in extant populations, the migrants en route seem to have settled in the Thar despite extremely harsh and resource-scarce conditions.^11,12^ The earliest evidence of farming, predominantly of wheat and barley, and early crafts like pottery could be traced to the excavations from Mehrgarh, the Neolithic sites (c. 7000 BCE) of South Asia^13^ that share boundaries with the northwestern regions of Thar. These practices flourished during the IVC from 3300 to 1300 BCE but declined due to climatic changes around 1900 BCE.^14^ Populations subsequently migrated eastwards and settled in the Gangetic Plains, Gujarat, and further southward coastal regions. ^11,12,15–21^ A few genetic studies suggest that modern-day South Asian populations are primarily a mixture of two major ancestral components: Ancestral North Indian (ANI) and Ancestral South Indian (ASI).^20–22^ However, the ANI-ASI cline does not fully explain the subcontinental genetic diversity. This is evident from earlier studies of the Indian Genome Variation Consortium (IGV) that provided the first comprehensive genetic landscape encompassing extensive linguistic, geographic, ethnic, and socio-cultural boundaries within India and the work of many others.^23,24^

The Thar Desert’s rich tapestry, woven from its ancient past and vibrant present, reflects a composite of ancestries shaped by early settlements, successive waves of migration, and diverse cultural and craft practices preserved within today’s distinct endogamous communities. The Marwari dialect spoken across Rajasthan also reflects the linguistic spread of the Indo-European linguistic family.^25,26^ Mass migrations of people in the last century could also have contributed significantly to demographic and social impacts on regions in the western part of Rajasthan. In addition, the harsh environment of the Thar Desert, including high temperatures, aridity, frequent sandstorms, and scarcity of resources,^11,27^ may have together shaped human adaptation and population structure. Craft communities in the Thar, with their kinship-based structures, distinctive cultural practices, and histories of migration from diverse regions, may have further contributed to the unique genetic architecture of this desert population. This study aims to elucidate the genetic diversity of Thar populations, particularly in the background context of distinct craft communities. We report genetic insights into the peopling of the Thar from genome-wide data on 176 individuals spanning eight distinct occupational communities in four underexplored districts. By establishing a genomic baseline from the Thar, this study offers potential for risk stratification in native populations while augmenting efforts to uncover the genetic and cultural basis of adaptation to extreme desert environments.

## Materials and Methods

### Ethnography of the native population from the Thar Ecoregion

Our study focuses on the craftsmen of the northwest provinces of the Indian subcontinent. This region spans across four districts: Jaisalmer, Jodhpur, Barmer, and Bikaner. The populations native to these regions practice traditional crafts and art forms that include pottery, terracotta, woodcraft and carving, gold embossing and painting, leather craft (mojri), textile and embroidery, tie and dye, and folk dance (Table S1). Clusters of households from locations of artisan families representing different crafts and semi-nomadic groups engaged in pastoral and artistic traditions were documented for the study. All interactions with participants were in accordance with the ICMR National Ethical Guidelines for Biomedical and Health Research Involving Human Participants (2017).^28^ Ethical approval was obtained from the Institutional Ethics Committee of the Indian Institute of Technology Jodhpur. Informed consent was obtained from all participants. Population identifiers as well as sample names were anonymized to safeguard their privacy and prevent stigmatization risk of isolated and endogamous communities.

### Identification of populations and sampling

Based on the ethnographic study, we identified eight native communities and one migrant community in the Thar Desert region. These communities are contrasting and diverse in terms of occupation, art forms, and sociocultural practices and are together referred to as Thar (TH) populations . A convention was evolved to label the population according to their locality and community practices in the samples. The population names were labelled as TH-IJE-X-(R/NR/MU/SN/U/PR), where TH-IJE indicates the study ID from the IIT Jodhpur ethnography study (TH-IJE) followed by the serial number of the sampling population (X) and from Rural (R), Nomadic (N), MU (Migratory Urban), SN (Semi-Nomadic), U (Urban), and Pastoralist (PR) (Figure 1) (Tables S1, S2).

**Figure 1.**
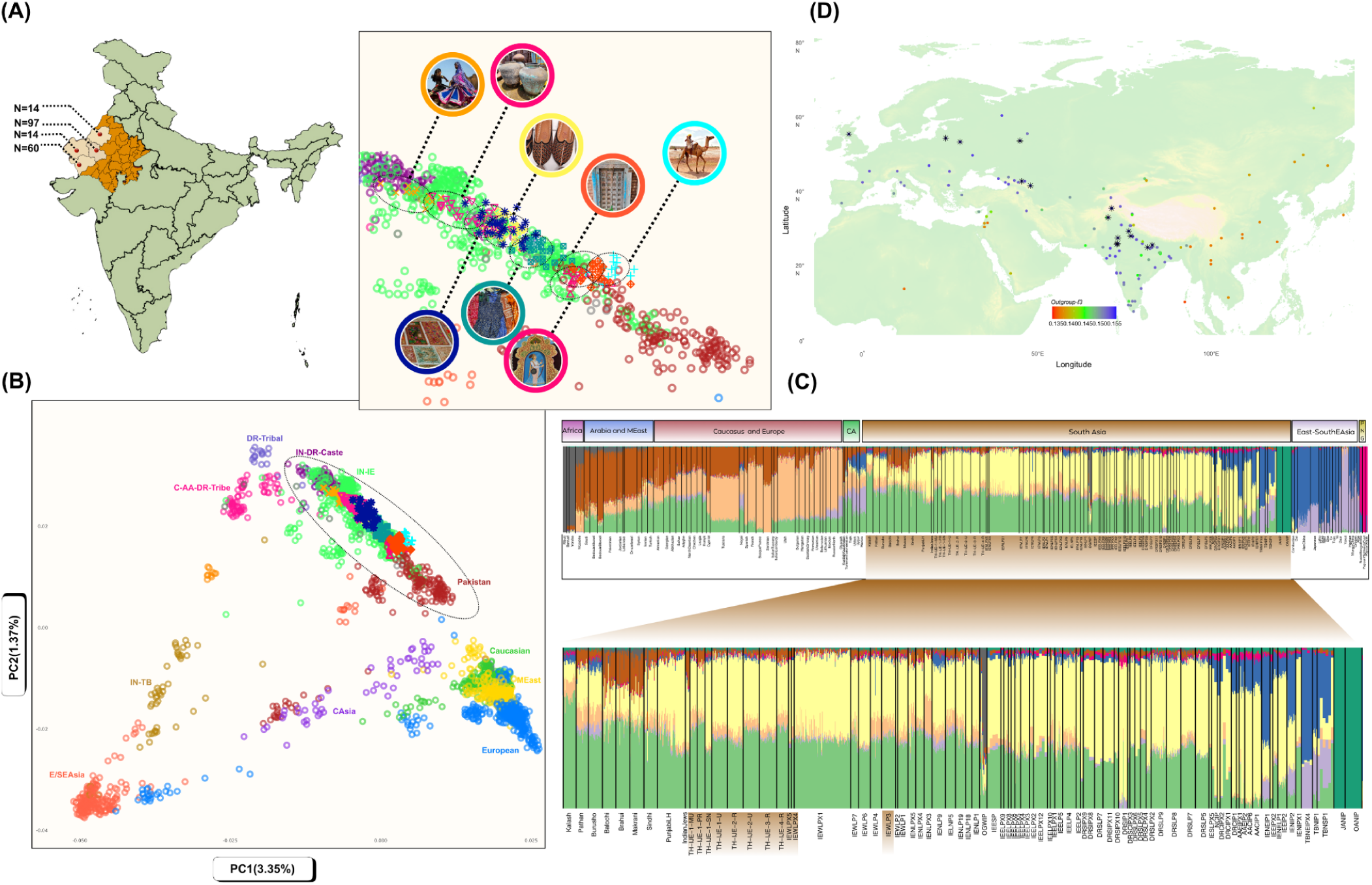
Genetic structure and affinities of populations from the Thar Desert, Rajasthan. (A) Map showing the geographical and sampling locations of the newly studied population groups from four major districts of Rajasthan (Thar Desert region of India). (B) Principal component analysis (PCA) of the merged dataset with worldwide reference populations (see Methods). The zoomed inset highlights the study populations from representative crafts, illustrating their placement within the north–south genetic ancestry cline of South Asia. (C) Unsupervised clustering results from ADMIXTURE analysis at K = 9, displayed as a stacked bar plot. Global populations are ordered geographically (Africa, Arabia and the Middle East, Caucasus and Europe, Central Asia, Southeast and East Asia, Papua New Guinea, and South Asia). The zoomed inset focuses on South Asian populations, with study groups labelled (see Methods). (D) Outgroup f3 statistics of the form f3(TH-IJE-4-R, X; Yoruba), represented as a geographical gradient map. Populations with the highest allele sharing are indicated in blue (high affinity), green (moderate affinity), and red (low affinity) to West Eurasian. Asterisk (*) the top 15 populations showing the highest amount of shared drift with TH-IJE-4-R.

### Genotyping and Integration of Thar with global datasets

A total of 185 individuals were genotyped using the Infinium Global Screening Array 24+ v3.0 + multi-disease ∼700K markers following the manufacturer’s protocol. DNA was extracted from peripheral blood,^29^ and GenomeStudio v2 (Illumina) was used for initial QC based on gentrain score, call rate, and cluster separation.^30^ After QC, 176 samples were retained with ∼660K markers. Genotyping accuracy was improved using cluster files generated from the Indian reference dataset.^31^ To place Thar populations in a broader context, genotype data were merged with published datasets representing major continental and Indian populations.^23,32–40^ To harmonize the nomenclature with respect to geographic and linguistic affiliations of Indian populations, we followed a convention of renaming the populations based on the convention followed in the Indian Genome Variation Consortium.^35^ Details of population inclusion are provided in Table S2.

AADR-1240K panels from prior studies were incorporated to represent panel of variations from ancient sites reported in previous studies^22,41–44^ that identify multiple layers of ancient West Eurasian ancestry in South Asia. These include Mesolithic Eastern European Hunter-Gatherers (EEHG) and Caucasus Hunter-Gatherers (CHG), Neolithic Iranian farmers (∼9,000–7,000 years ago) from the Zagros, and Neolithic Anatolians. Steppe Early–Middle Bronze Age (EMBA) groups (Yamnaya-related) arose from EEHG and CHG. Additional ancestry sources include Neolithic/Bronze Age Iran-Turan, Siberian hunter-gatherers, Botai, Namazga, Bactria–Margiana Archaeological Complex (BMAC), Indus Periphery, Historical South Asia (SouthAsia_H), SPGT (likely Swat Protohistoric Grave Type), and India_Roopkund.

Data merging was performed using EIGENSOFT (convertf, mergeit).^45^ After applying QC filters included call rate, genotyping error (0.10), MAF (0.01), individual missingness (0.15), and HWE (1e−06), yielded 77,561 variants and 2,812 individuals (Dataset 1). After LD pruning in plink1.9 (--indep-pairwise 200 50 0.4), 71,414 variants remained (Dataset 2) for PCA and admixture analysis. A merged subset with the 1240K panel^46^ (Dataset 3) containing 85,393 variants was used to explore Thar’s genetic affinities with ancient and modern Eurasian and South Asian populations (Table S2).

### Genome-wide autosomal analysis

We conducted genome-wide autosomal analyses to investigate the genetic structure of Thar populations in relation to neighboring and reference groups. We performed PCA at the individual level for both aDNA Data and among modern populations after pruning SNPs using SMARTPCA from the EIGENSOFT package,^45^ with default parameters (numoutlieriter: 0, lsqproject: Yes). Results were visualized in R using the *ggplot2* package.^47,48^ ADMIXTURE^49^ analysis was carried out after pruning SNPs in strong linkage disequilibrium with Plink v1.9.^50^ We also carried out 5-fold cross-validation, varying the number of ancestral populations from K = 2 to K = 18, each with 25 replicates using different random seeds. The optimal K value was chosen based on the lowest cross-validation error and highest log-likelihood. Pairwise F_st_ values were estimated between Thar populations and neighboring groups using SMARTPCA in EIGENSOFT^45^, with the parameters F_st_ only: YES and phylip^51^ outname enabled. Block jackknife resampling was used to calculate standard errors.

In order to understand the relationships between populations in terms of common ancestry, genetic drift, and gene flow, we estimated a maximum-likelihood tree using TreeMix v1.13^52^ based on genome-wide allele frequency data from Indian populations, where OANIP (Andamanese) was used as an outgroup to root the output tree based on the ML graph visualizing population splits and admixture events without any migration edges in order to provide a simple graph tree.

As ADMIXTURE is not a formal test for gene flow between populations, we conducted three-population admixture tests (f3 statistics)^53^ to assess shared genetic drift and evolutionary history. In pairwise Outgroup f3, we considered Thar and its most adjoining region population (Thar/Pakistan/IEWLPX1, Pop2; Outgroup) test, where Pop2 are all the other neighboring populations, such as South Asians, Europeans, Middle Easterners and Caucasians from Dataset 3, and Pop2 as outgroup African populations (Yoruba). We inferred the extent of admixture and gene flow with neighboring populations using the threshold significant value of the Z-score -2 to +2 range with jackknife standard errors for the test population.

We also employed f4 statistics using the qpdstat program in Admixtool 7.0.2^53,20^ with many combinations to assess gene flow and investigate derived allele sharing between Thar and other populations. The proportion of ancestral North India was also estimated from the f4 ratio estimation using AdmixTools 7.0.2,^53,20^ which analyses the correlation of allele frequency patterns to determine the proportion of Ancestral North Indian (ANI) ancestry in different Indian ethnic groups. The ANI ancestry proportion was calculated based on the model outlined by Moorjani et al.^20^ To assess the uncertainty associated with ANI ancestry proportions we used two distinct geographical extremes: the Abkhazian and Basque populations reported earlier.^20^

We used qpWave and qpAdm,^53^ using the admixr^54^ package from R, to estimate ancestry proportions in South Asians by modelling them as a mixture of ‘reference’ populations while utilising shared genetic drift with a set of ‘outgroup’ populations. In this analysis, the group OANIPa* was taken as a proxy for the South Asian component, representing Ancient Hunter-Gatherers (AHG), since they lack recent West Eurasian ancestry.^55^ Using the 1240K-merged dataset, the test population was modeled with three reference groups: OANIPa* (n = 17; non-African Andamanese islanders), Mesolithic and Late Bronze Age Steppe-related groups (Central/Western Steppe-MLBA), and Indus Periphery/Diaspora (Indus Periphery) as the proximal model, while Namazga_CA was included as the distal model. To assess mixture proportions, we used a diverse set of modern and ancient populations; for distal, we used outgroups such as Ethiopia_4500BP, SG, MA1, PPN, EEHG, WEHG, Natufian, ESHG, and Dai. For proximal, we used Ethiopia_4500BP. SG, Anatolia_N, PPN, EEHG, WEHG, Sarazm_EN, ESHG, and Dai, including the fitted model as right outgroup populations, We also used Sarazm_EN as a representative of the Neolithic Iranian-related source, with the following outgroups in the fitted model such as Ethiopia_4500BP, WEHG, EEHG, Ganj_Dareh_N, Anatolia_N, ESHG, Dai, Russia_Samara_EBA_Yamnaya, and Vietnam_BA. followed using Narasimhan et al., Kerdoncuff et al. ^22,56^ allowed us to explore the genetic structure of Thar natives and their ancestral affinities with different prehistoric populations.

We computed weighted linkage disequilibrium (LD) statistics using ALDER^57^ to estimate the timing of admixture events in the Thar population. The method relies on the exponential decay of LD generated by admixture, with the decay rate providing an estimate of the time since admixture. To improve accuracy, the algorithm measures SNP correlations in the admixed population and applies weights based on allele frequency differences between reference populations. This approach enhances admixture-related LD signals while reducing the influence of background LD, thereby minimizing potential biases. For our analysis, we used West Eurasian and AASI populations as references. Standard errors were calculated using chromosome-based jackknifing, and a generation time of ∼30 years was assumed for converting results into calendar years. We then used ASCEND v10.1.1,^58^ which measures allele sharing correlation between pairs of individuals across the genome to infer the age and strength of the founder event in the Thar population. We used a merged dataset with a 1,240k panel dataset,^46^ which includes ∼138k SNPs. Mbuti samples were considered as the outgroup. The founder age was calculated assuming a generation time of 30 years.

For inferring demographic history and estimating the effective population, we used HapNe^59^ based on LD summary statistics and composite likelihood given in the default parameters. Furthermore, to identify runs of homozygosity (ROHs) across the genome, we used PLINK v1.9.^50^ with the --homozyg function. Each window spanned 1000 kb and contained at least 100 SNPs. To account for possible genotyping errors and missing data, up to one heterozygous and five missing calls were allowed per window. Only ROHs ≥1000 kb were retained for analysis. Groups with fewer than four individuals were excluded to ensure reliable population-level estimates. For each population, we calculated the mean number and average length of ROH segments. Results were visualized in R using the ggplot2 package.^48^

We phased genome-wide dense SNP data using SHAPEIT2^60^ and used the Refined IBD and MergeIBD algorithms^61^ for identity by descent (IBD) analysis with default settings on a phased dataset. Segments greater than 1 cM were analyzed. Estimations were made on an individual basis, and the average IBD segment per pair of individuals was calculated by dividing the total number of segments by the possible combinations of pairs within each population. For IBD score within the population, we used the sample size for normalisation, and computed the raw IBD score only for the total length of IBD segments between 3 and 20 cM and used as mentioned in Nakatsuka et al 2017.^62^

The phased genome-wide dense SNP data were used for ChromopainterV2 to estimate the coancestry matrix, followed by running FineSTRUCTURE^63^ to perform Bayesian clustering and identify fine-scale population structure, and finally, the results were interpreted using the generated dendrogram and clustering assignments. We used ChromoPainterV2 and ran the analysis twice. First, we performed an initial run on a subset comprising 1/10 of the population (one individual per population), using 10 EM iterations. From this, we took the weighted average of the -n (mutation rate) and -m (switch rate) parameters to fix the mutation and switch rates for the full analysis. Specifically, we used the values inferred from a subset of representative chromosomes (1, 3, 7, 10, and 22) as described in Montinaro et al^64^ which resulted in -n = 555.757569 and -m = 0.000337 Using these inferred values, we then ran the analysis on all chromosomes with all individuals, applying an all-to-all donor strategy. The results of the mean chunks heatmap were visualised using the ggplot2 package in the R software.^48^

### Uniparental lineage analysis

We analyzed maternal lineages using mtDNA from 176 individuals with the rCRS as a reference. mtDNA haplogroups were assigned with HaploGrep3,^65^ based on PhyloTree Build 17 Forensic Update 1.2.^65–67^ Y-chromosome haplogroups were determined using Y-LineageTracker^67^ for the 63 male individuals out of the 176, following the ISOGG Y-DNA phylogenetic tree (2019–2020).

## Results

### Thar desert populations along the global genetic cline

We studied the genetic landscape across the Thar Desert in northwestern India, within the broader context of South Asian and Eurasian groups. The study populations were carefully selected based on their traditional crafts and cultural practices that are maintained within distinct endogamous, kinship-based groups from rural, urban, and seminomadic settings (Table S1). This included the Thar Urban leather artisans (TH-IJE-1-U), tie-dye artisans (TH-IJE-2-U), and the Migrant gold embossing artists (TH-IJE-MU); the Pastoralist from Rural region TH-IJE-1-PR; the Semi-Nomad/Performing Artist (TH-IJE-1-SN); potters (TH-IJE-2-R), embroiderers and weavers (TH-IJE-3-R), and wood carvers/carpenters (TH-IJE-4-R). This approach not only provided authenticity but also allowed us to directly assess the correspondence between genetic structure and craft as well as culture-based genetic affiliations and origins.^1–3,5,6^

Principal component analysis (PCA) with a variance of 3.35% PC1 and 1.27% PC2 reveals a genetic gradient extending from West Eurasia through South Asia to East Asia. The Thar craft populations occupy an intermediate position between groups along the western frontier and certain Indo-European–speaking populations from northwestern India, forming part of a north–south genetic cline (Figures 1, S2). This pattern of affinity is further supported by F_st_ values (Table S3, Figure S1), which indicate close genetic relationships between the Thar populations, western frontier groups, and populations from northern and western India. Nevertheless, some Thar groups deviate from this continuum, suggesting genetic differentiation within the region. Consistent results were observed in both ADMIXTURE (Figures 1, S3, Table S4) and F_st_ analyses (Figure S1, Table S3). In particular, the pastoralist (TH-IJE-1-PR), migrant artists (TH-IJE-1-MU), and wooden craft communities (TH-IJE-4-R) display closer genetic affinities with Pathan and Sindhi populations, as well as Indo-European–speaking groups from the Gangetic Plains and Northwestern India (Figures S3 and S9). By contrast, the potter community (TH-IJE-2-R) and the seminomadic group of traditional folk dancers (TH-IJE-1-SN) are distantly positioned at the lower end of the north–south genetic cline.

ADMIXTURE analysis (based on the lowest CV error value and log likelihood), at K=9 (Figure S3 and Table S4), reveals further that the genetic structure of Thar populations reflects variable contributions from South Asian, Middle Eastern, and European ancestries. The northwestern populations share major ancestry components with several South Asian groups (colors—yellow and green in Figure S3). The green component is highest in the Kalash population and is widely distributed across Central Asian, Middle Eastern, and European groups. Both TH-IJE-1-MU and TH-IJE-4-R harbor comparable levels of the European component (brown), consistent with their proximity to Indo-European–speaking populations of the gangetic plains and the northwestern frontier. This corroborates the observations from the admixture analysis (Figure 1). TH-IJE-1-SN shows a higher proportion of Ancestral South Indian (ASI)-related ancestry, similar to the isolated Dravidian-speaking population DRSIP1, whereas the Kalash population displays the lowest levels of ASI but the highest levels of Ancestral North Indian (ANI)-related ancestry. These observations are consistent with historical interactions along the northwestern frontier.

The population relationships and demographic events reconstructed using f₃ statistics that provide formal tests for admixture and shared genetic drift among populations reveal an interesting pattern. Outgroup f3 statistics for genetic drift with Yoruba as the outgroup reveal that the Thar subgroups—TH-IJE-4-R, TH-IJE-1-PR, and TH-IJE-1-MU, with some other Indo-European-speaking populations, exhibit elevated levels of shared genetic drift with West Eurasian populations. The strongest signals are observed with the Kalash, Pathan, Lezgin, Chechen, Belarusian, Lithuanian, and Scottish Orkney groups (z-score > 100) (Table S5, Figure 2), suggesting a clear affinity between these Thar groups, West Eurasians, and populations located along the northwestern frontier. In contrast, other subgroups like TH-IJE-1-SN and TH-IJE-2-R showed stronger levels of shared drift with Indian populations from the Gangetic plain Indo-European cline and Dravidian-speaking South Indian caste groups. Notably, TH-IJE-1-SN follows a drift-sharing pattern resembling that of Gujarati populations. These findings from the outgroup f3 analysis are further corroborated by D-statistics using qpDstat (Tables S6). Specifically, the Pathan and Kalash populations exhibit significantly higher levels of gene flow with the Thar individuals compared to most other North Indian groups. Contrastingly, TH-IJE-1-SN does not display significant allele sharing with West Eurasian populations when European and Pakistani populations, such as French and Georgians, Iranians, Pathans, Balochi and Kalash, were taken as reference populations (Table S6, Figure 3).

**Figure 2:**
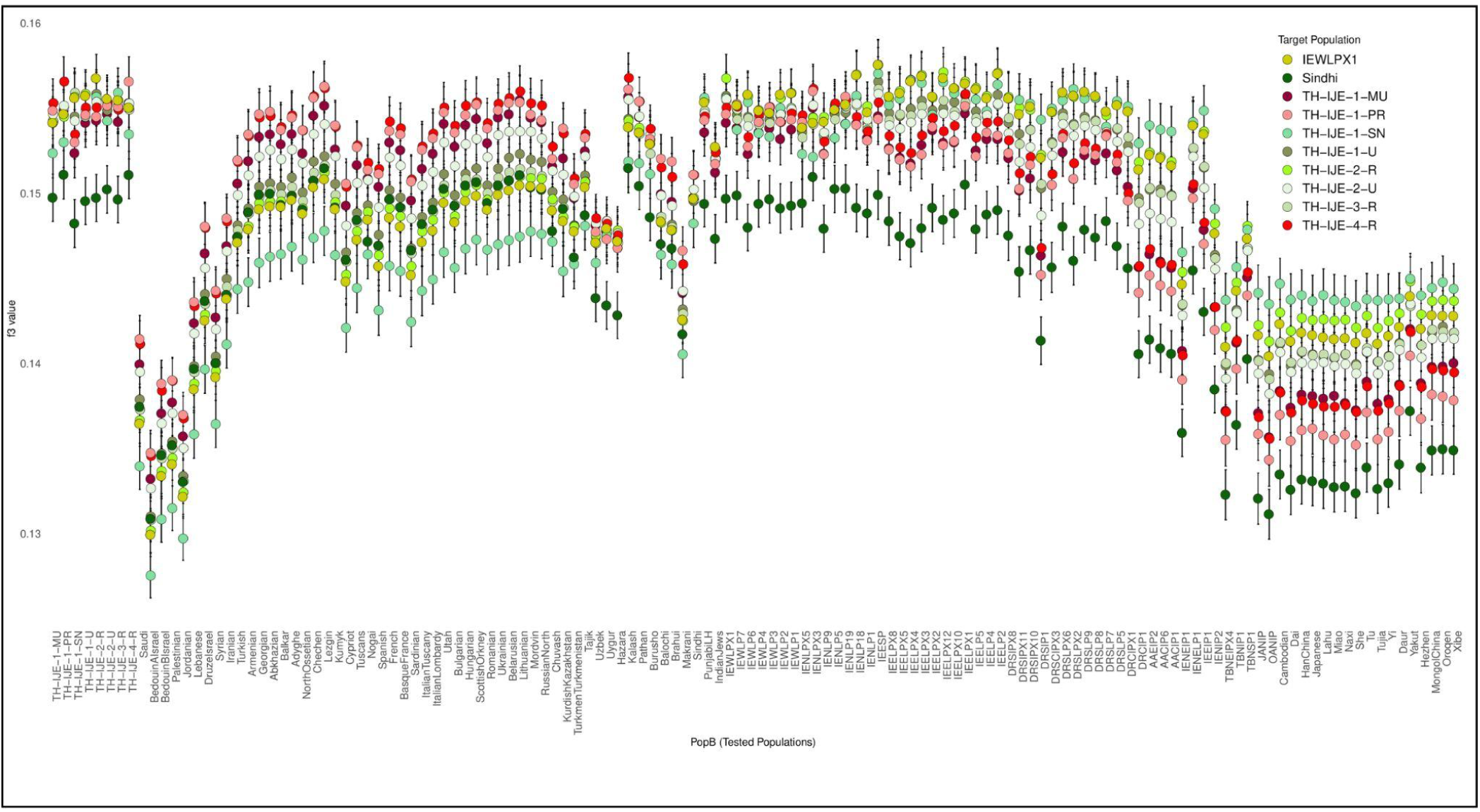
Derived allele sharing of modern Thar Desert populations with other Eurasian groups. Scatter plots show the results of the outgroup f₃ test of the form f₃(Thar/IEWLPX1/western frontier population, X; Yoruba), where X represents any other modern Eurasian population. Error bars indicate jackknife-derived standard errors, highlighting the relative allele sharing between the study populations showing higher shared drift compared to IEWLPX1 and Sindhi population towards Western Eurasian groups.

**Figure 3:**
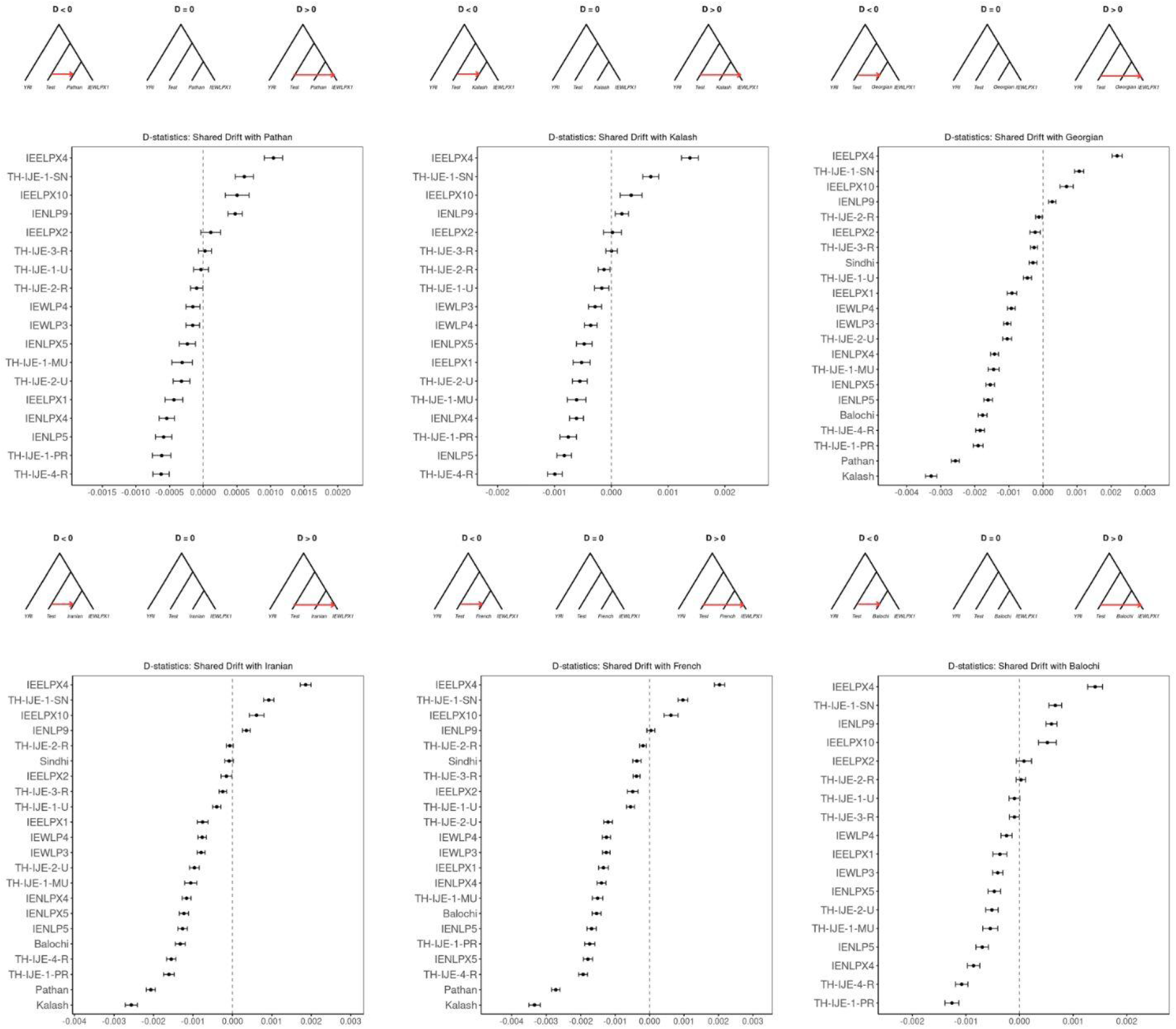
Gene flow and relative affinities of Thar Desert populations with Indo-European groups and West Eurasian. D-statistics in the form D(pop1, Yoruba; pop2, pop3) were computed, where pop1 corresponds to the Thar individual IEWLPX1, pop2 represents modern South Asian populations, and pop3 includes South Asian or West Eurasian reference groups. The results highlight patterns of allele sharing among the western frontier populations and Indo-European groups of North/West Indians. The Thar craft communities show particular genetic proximity to Pathan and Kalash populations, and demonstrate their strongest affinities with western frontier and european groups compared to other South Asians

We also computed f3-statistics to evaluate the extent of formal admixture of Thar populations along with groups from the Gangetic Plains and Rajasthan with Ancestral North Indian and Ancestral South Indian lineages. Ancient and modern West Eurasian populations were used as proxies for the ANI component and DR-S-IP1 for the ASI component. We found that populations such as TH-IJE-1-MU, TH-IJE-1-U, TH-IJE-2-U, and TH-IJE-1-SN did not yield significantly negative f3-values, suggesting a stronger affinity toward a single ancestral source and limited evidence of direct ANI–ASI admixture. By contrast, groups including TH-IJE-4-R, IEWLP3, TH-IJE-3-R, and TH-IJE-1-PR from the Thar region exhibit significantly negative f3-values, consistent with admixture signals observed among Gangetic Plains populations and northwestern Indo-European–speaking groups (Table S7). These results suggest that while certain Thar populations retain genetic continuity with one predominant ancestry, others share admixture patterns characteristic of broader ANI–ASI interactions that have shaped the genetic landscape of North India.

To further estimate the proportion of the Ancestral North Indian (ANI) component, we applied the f4-ratio method of ADMIXTOOLS. Higher proportions of ANI ancestry were observed in populations such as the Kalash, Pathan, and northwestern groups IENLPX3 (Ror), IENLPX4 (Gujjar), and IENLPX5 (Kambhoj). Among the Thar groups, TH-IJE-4-R, TH-IJE-1-PR, and TH-IJE-1-MU also exhibit elevated ANI ancestry compared to most other South Asian populations. In contrast, TH-IJE-1-SN displayed a notably reduced proportion of shared ANI ancestry (Table S8). Consistent patterns are observed in the maximum likelihood tree of TREEMIX, which estimates population-specific drift and divergence. Thar populations cluster within the clade of Indian Indo-European caste groups. Compared to other Thar populations, TH-IJE-1-MU and TH-IJE-2-U exhibit higher levels of population-specific drift, as reflected by longer branch lengths relative to most South Asian populations, except Kalash. Furthermore, several northwestern subcontinental populations were positioned within the same clade as TH-IJE-1-MU, TH-IJE-1-PR, and TH-IJE-4-R. In contrast, TH-IJE-1-SN consistently remains more distantly related to this clade (Figure S5).

Analysis of identity-by-descent (IBD) using RefinedIBD patterns shows a reduction in sharing in a clinal gradient, declining westward from groups near the northwestern frontier and northward in the Indo-European groups of North India. This decline was even more pronounced with Dravidian and Austroasiatic groups from southern and eastern India, respectively. Notably, the migrant artist group from the western frontier (TH-IJE-1-MU) exhibited elevated IBD sharing with European populations, a pattern also observed in the Pathan population. Similarly, TH-IJE-1-PR and TH-IJE-4-R show increased IBD sharing with neighboring northwestern frontier populations such as Balochi and Sindhi, as well as with Indo-European groups from the northwest and Gangetic Plains, and to a lesser extent, with Dravidian populations (Figure S6). Within-population IBD analyses normalized by population size,^61,62^ we observe that individuals of the Thar population, such as TH-IJE-1-MU, TH-IJE-1-U, and TH-IJE-2-U, exhibit the highest IBD scores among the Indian populations and other craft artisans were also showing higher than that of Finns (Table S9). This finding corroborates earlier accounts of increased inbreeding and strong genetic drift. Elevated levels of homozygosity were also observed in most artisan communities of the Thar Desert. Both the mean number and mean length of runs of homozygosity (ROH) segments >1 Mb were significantly higher compared to other Indian groups. These findings indicate reduced genetic diversity and recent population isolation (Figure S7).

To investigate the demographic history of the Thar populations, we applied complementary methods focusing on founder events and admixture timing. Using the ASCEND (Allele Sharing Correlation for Estimating Demography) algorithm, we inferred founder intensities ranging from 2.0% to 4.6% across Thar subgroups, indicating varying degrees of population bottlenecks and long-term endogamy (Figure S8). Application of the HapNe-LD approach further identified recent bottleneck events in groups such as TH-IJE-2U, TH-IJE-1-MU, and TH-IJE-2R, with estimated timings within the last 400–500 years (Figure S8). These findings suggest that certain Thar subgroups experienced relatively recent reductions in effective population size, likely linked to social isolation and strict endogamy. Furthermore, for the timing of West Eurasian admixture, we applied weighted linkage disequilibrium (LD)-based inference using ALDER. Admixture dates were successfully estimated by modelling the Thar populations as mixtures of two reference sources: modern West Eurasian groups and South Asian groups. Following earlier studies, Austroasiatic-speaking groups were used as proxies for AASI ancestry to represent the South Asian source. Under this, we inferred admixture events in Thar artisan populations, with most groups showing evidence of West Eurasian ancestry dating to approximately 60–80 generations before present. Notably, TH-IJE-2U exhibits a more recent admixture event, dated to ∼30 generations before present (Table S10).

Complementary insights were obtained from ChromoPainter and finestructure analysis based on haplotype chunk counts. Individuals from TH-IJE-4-R, TH-IJE-1-PR, and TH-IJE-1-MU share a greater mean number of haplotype chunks with European and Caucasian populations compared to other Thar artisan groups. By contrast, TH-IJE-1-SN exhibits the lowest levels of haplotype sharing with these groups (Figure S9). Collectively, these results reveal a gradient of westward genetic affinity, with most Thar populations showing stronger haplotype sharing with populations along the northwestern frontier than with other South Asian groups, including Dravidian- and Austroasiatic-speaking communities.

### Determination of ancestral proportions in Thar populations in a global context

Using PCA, outgroup-f₃, and D-statistics analyses, we found that the Thar populations exhibit stronger genetic affinity with EMBA groups than with later Steppe MLBA groups. Within the Thar, individuals from TH-IJE-4-R, TH-IJE-1-PR, TH-IJE-1-MU, and TH-IJE-2-U cluster more closely with proximal sources such as South_Asia-H, Indus Periphery, SPGT, Namazga Chalcolithic, and BMAC, distinguishing them from other Thar subgroups with higher indigenous components that cluster farther along the South Asian cline (Figure S4).

Outgroup-f₃ analyses in the form of f(test, ancient pop; Yoruba) showed that these Thar groups shared elevated genetic drift with ancient reference populations, including EEHG, CHG, Neolithic Iranian farmers, and Neolithic Anatolian, compared to other South Asian groups (Figure 4, Table S11).

**Figure 4.**
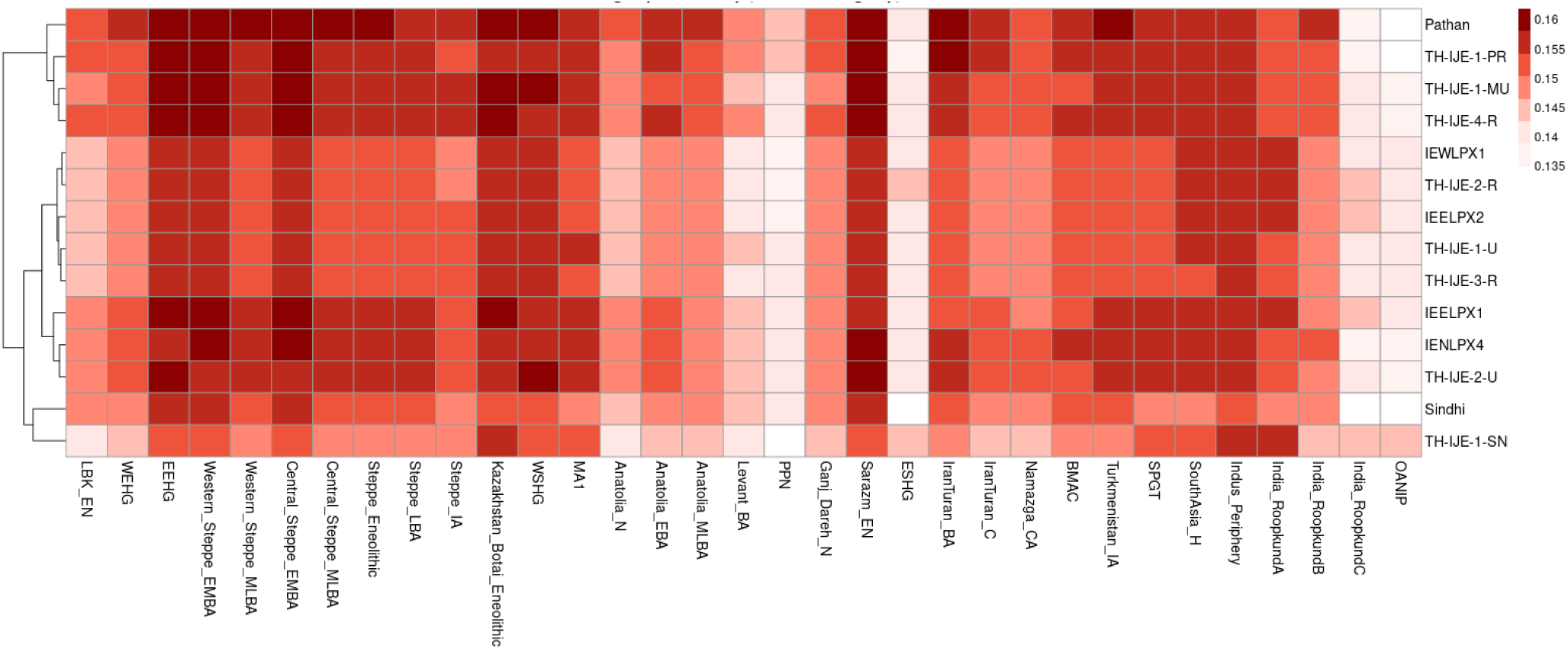
Outgroup-*f*₃*-*based genetic affinity heatmap across ancient Eurasian populations. The heatmap shows shared genetic drift values (f₃[test, ancient population; Yoruba]) between the Thar and other South Asian populations with a range of ancient Eurasian groups. Broad geographic and temporal categories include European and Steppe-related, Near Eastern and Iranian, and South and Central Asian populations. Higher f₃ values in darker color in the Some of the Thar groups indicate stronger shared genetic drift and closer genetic affinity with these ancient sources.

We applied D-statistics in several configurations to explore allele sharing with ancient populations. Using D(test, Yoruba, AncientPop1, AncientPop2), we assessed affinities between South Asian groups and ancient sources. Across most South Asian populations, including the Thar groups, the D-statistics reveal broadly similar patterns, where Western Steppe_EMBA consistently shows stronger affinity compared to Central and Western Steppe_MLBA and Namazga_CA (Table S12). Additionally, we also observed stronger affinity toward Sarazm_EN compared to Ganj_Dareh_N across all South Asian populations, suggesting that ancestry related to Sarazm_EN may better represent the Iranian farmer–derived component. The Thar populations show overall affinity with Indus_Periphery, comparable to that seen in other Indian groups from the south. However, TH-IJE-1-PR, TH-IJE-4-R, and TH-IJE-1-MU display similar allele sharing and affinities with both Middle and Late Bronze Age Steppe groups and Indus_Periphery. These corroborate earlier observations on northern and western Indian populations (Table S12).^22,68^

When we used more proximal sources evident from PCA, we found TH-IJE-1-PR, TH-IJE-1-MU, and TH-IJE-4-R show comparable levels of allele sharing with Indus Periphery populations when tested with BMAC, Namazga_CA, and SPGT. In contrast, among Indian populations, central Austroasiatic and Dravidian groups appear closer to the Indus Periphery, South_Asia-H, or SPGT (Table S13). Notably, TH-IJE-1-PR exhibits a particularly strong affinity with BMAC compared to other Indian populations. Interestingly, when compared to present-day Iranians, TH-IJE-1-PR, TH-IJE-4-R, and TH-IJE-1-MU show a stronger affinity with EEHG. This reflects a more ancient admixture with the populations around this region and gene flow from a Central Asian or Steppe-related West Eurasian source (Table S13). Both TH-IJE-1-MU and TH-IJE-1-PR also display significant allele sharing with Turkmenistan_IA compared to SPGT, highlighting Central Asian links in their genetic profiles (Table S12).

To explore southern ancestry components, we compared affinities with the ASI proxy (DRSIP1) relative to SPGT and South_Asia-H. Across South Asian populations, TH-IJE-1-PR, TH-IJE-4-R, and TH-IJE-1-MU show higher affinity with SPGT and South_Asia-H, similar to other western frontier populations, whereas populations from central and southern India exhibit stronger affinity toward the ASI proxy (Table S12) . To further elucidate the ancestral affinity, we implemented D(test, TH-IJE-[1PR/4R], AncientPop, Yoruba), where either of the three Thar populations was used as a target with the test as other South Asian populations. We found that among the northwestern groups, TH-IJE-1-PR and TH-IJE-4-R show more affinity with Chalcolithic and Neolithic populations such as Namazga_CA and Ganj_Dareh_N, as well as with historical groups like SPGT, South_Asia-H, and BMAC. Additionally, with other ancient sources, including CHG, EEHG, Anatolia_N, and other steppe-related populations, these show affinity patterns comparable to those observed in Pathan and Kalash groups from South Asia (Table S13).

Admixture modelling using f₄ ratios by a model of f₄(Mbuti, EEHG; Test, non-African Andamases) / f₄(Mbuti, EEHG; Central_Steppe_MLBA, non-African Andamases) further supported these findings. We detected substantial Central Steppe MLBA ancestry (proxy for ANI-related ancestry when using Namazga CA / Indus periphery and AHG for a better fit model) in the Thar, ranging from 48.7 ± 1.9% to 34.6 ± 2.2%. This proportion was lowest in TH-IJE-1-SN and TH-IJE-2-R but higher in Pathan, Kalash, and Thar groups such as TH-IJE-1-PR and TH-IJE-4-R (Table S14).

Using qpAdm^44,53^ three-way admixture models (Western/Central Steppe MLBA, Indus Periphery cline, and AHG/non-African Andamases), we found that many populations fit the qpAdm model (p > 0.05), though some failed, suggesting that this framework may not fully capture the complexities of ancestries along the north–south cline. These estimated results show a consistent proportion, as mentioned in previously studied populations. Notably, TH-IJE-1-PR seems to have a predominantly ancient Iranian farmer-related proportion with minimal AHG contribution, closely resembling the Kalash profile. By contrast, TH-IJE-1-SN shares genetic proportions that are similar to the Dravidian-speaking caste group (Tables S15, S16) (Figure 5).

**Figure 5.**
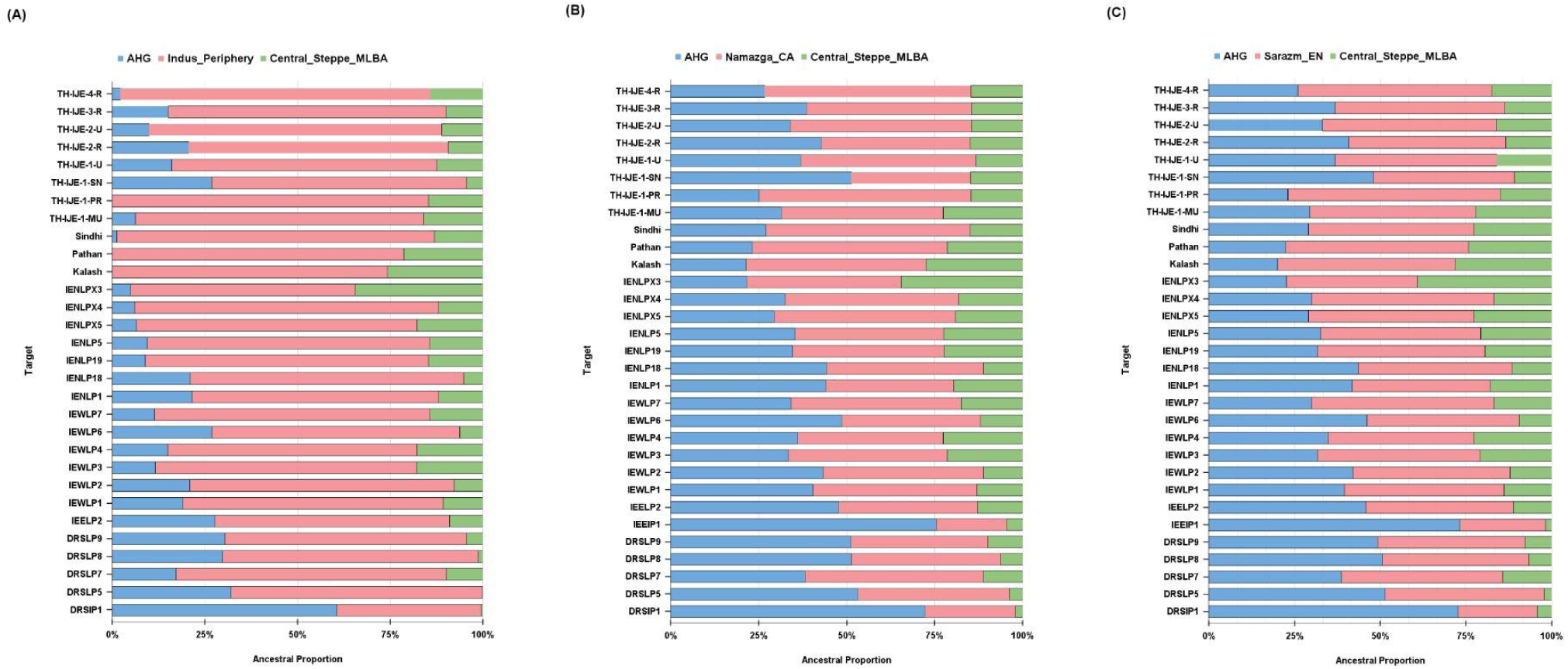
Admixture modeling showing the proportions of ancient West Eurasian ancestry in the Thar artisan populations. (A) **Proximal model** using OANIP (AHG), *Indus_Periphery*, and *Central Steppe MLBA (proxy for ANI)* as source populations to infer recent ancestry contributions. (B) **Distal model** using OANIP (AHG), *Namazga_CA*, and *Central Steppe MLBA* as ancestral sources representing Iranian-related and Steppe ancestries. (C) **Alternative distal model** using *Sarazm_EN* to test the contribution from Neolithic Iranian-related ancestry from the Iran–Turan region. Each color denotes a distinct ancestral source, and represents the fraction of total ancestry estimated for each population.

Among the Thar individuals, TH-IJE-1-MU consistently shows the highest proportion of Steppe ancestry, followed by other Thar groups and northern/western Indian populations such as IENLPX4, IENLP5, and IEWLP4. The Steppe component in these groups is comparable to that observed in Pathan and Kalash, while TH-IJE-4-R and TH-IJE-1-PR display relatively higher proportions of Iranian-related ancestry (Tables S15, S16). Overall, the ancestry proportions in the Thar artisan groups fall within the range observed across other South Asian populations. In the distal admixture model, most populations showed a good fit (p > 0.05). Consistently, both distal and proximal models indicate qualitatively higher Bronze Age Steppe ancestry in TH-IJE-1-MU (Tables S15, S16). Based on results from outgroup-f3 and D-statistics, we also model by using Sarazm_EN as a proxy for the Iranian farmer related source, together with AHG as the ASI proxy and Central Steppe_MLBA as the ANI proxy. The model provided a good fit for most South Asian and Thar populations. (Tables S16) (Figure 4).

Both distal and proximal qpAdm models reveal patterns consistent with previously studied reconstructions of South Asian population history and episodes of population movement from the Steppe and Central Asia into South Asia during the Bronze and Iron Ages, while also underscoring the unique position of Thar subgroups at the intersection of cultural and demographic exchanges across the Indo-Iranian frontier.

### Gene flows mirror the genetic route of migrations from Europe to the Thar

To assess whether present-day Thar populations carry genetic signatures that corroborate the insights from ancient migration and adaptation histories, we examined the distribution of functional alleles underlying traits such as lactase persistence and skin pigmentation. We investigated the distribution of the lactase persistence-associated rs4988235 T allele (−13910*T) of an LCT gene that enables the digestion of milk into adulthood. Our analysis revealed marked heterogeneity in its frequency among populations of the Thar. Consistent with broader South Asian patterns, the T allele generally declines in frequency from North to South and from West to East. However, within the Thar populations, we observed distinct local enrichments. Compared to the previously studied Ror (IENLPX3) population,^68,69^ which shows the highest frequency (71.4%), the groups from our study exhibit extensive heterogeneity, with the pastoralist populations having the highest frequency of TH-IJE-1-PR (67.9%), followed by TH-IJE-4-R (40.5%), TH-IJE-3-R (29.0%), and TH-IJE-1-U (26.8%). Surprisingly, the least in TH-IJE-1-MU (Figure 6). The higher frequencies resemble those observed in European and West Eurasian pastoralist groups, where the allele is under strong positive selection due to reliance on dairy consumption. ^69–71^

**Figure 6:**
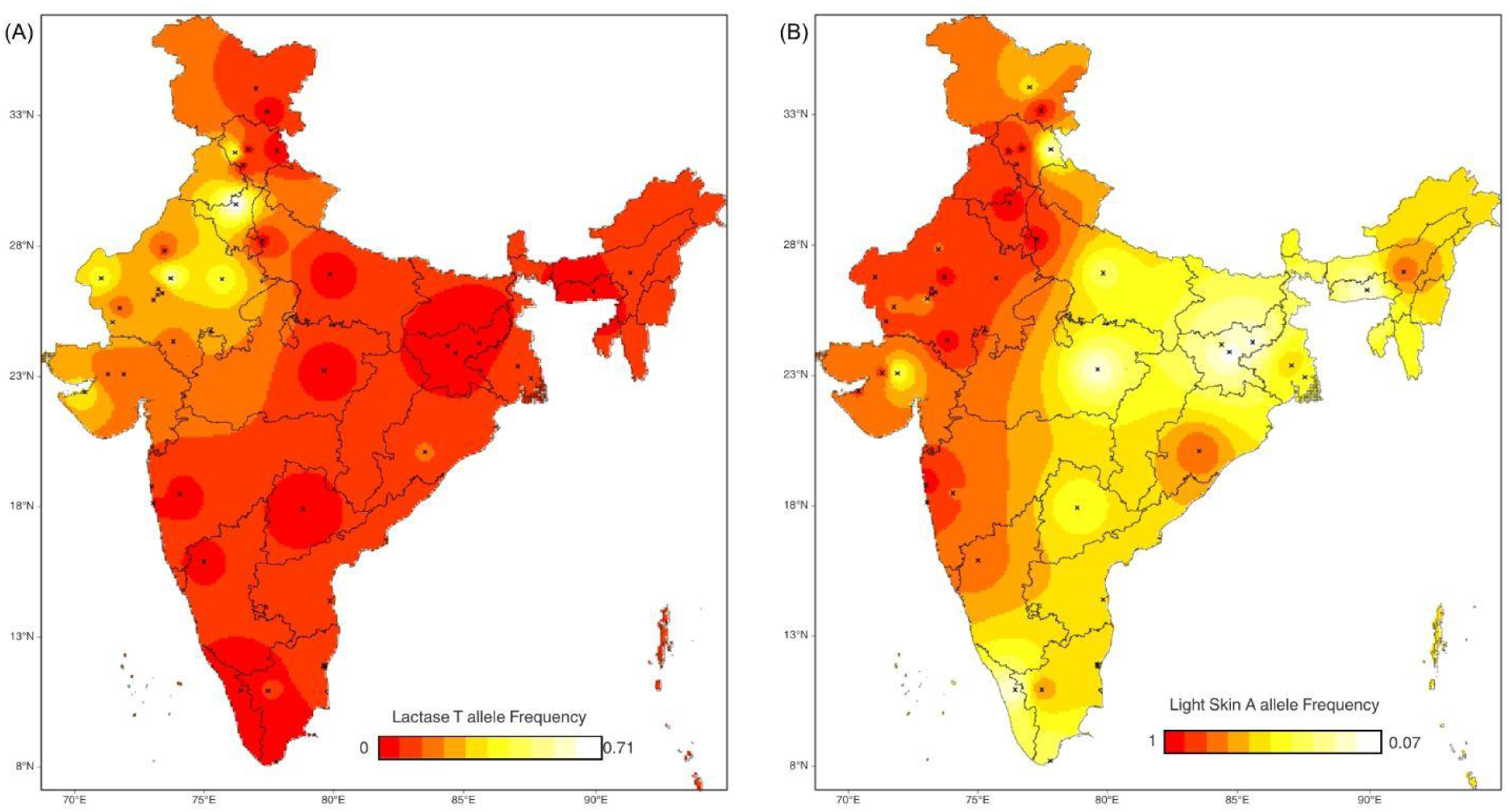
Spatial frequency map of India. Spatial frequency map showing allele frequency distribution for (A) LCT rs4988235 (T allele) and (B) SLC24A5 rs1426654 (A allele) across Indian populations. The color gradient represents the observed allele frequency range of the T allele of *LCT and* the A allele of *SLC24A5 in* North and Northwest Indian populations, showing similar frequencies as seen in European groups.

In addition, we also examined the rs1426654 variant in the SLC24A5 gene, a major contributor to lighter skin pigmentation in Eurasian populations.^72,73^ The derived A allele, which is nearly fixed in Europeans, also displays considerable frequency variation across the Thar region.^69–73^ It is most frequent in the IENLPX3, IENLPX5, IENLP5, TH-IJE-1-U, and TH-IJE-1-PR groups and is also elevated in several other North Indian populations. Interestingly, some groups of these also show high frequencies of the lactase persistence allele, although the two traits vary independently, suggesting parallel but distinct histories of adaptation for dairy consumption and pigmentation. Conversely, groups such as TH-IJE-1-SN and TH-IJE-2-R exhibit intermediate A allele frequencies, similar to many Indo-European and tribal-affiliated populations (Figure 6).

### Uniparental Genetic Analysis

The uniparental markers of the Thar populations reveal a tapestry of ancestries, reflecting both South Asian roots and multiple waves of external influence.^74–80^ On the maternal side, mitochondrial analysis revealed 55 distinct sub-haplogroups encompassing 5 of the 12 major global haplogroups. Over half of the individuals (51.1%) belonged to haplogroup M, the most widespread and ancient lineage of South Asia, while nearly one-third (31.8%) carried haplogroup U. However, haplogroups U7, W, and R2, which are associated with West Eurasian ancestry, are also consistently present in the three Thar populations studied. Smaller fractions of haplogroup H (5.7%) and the presence of lineages such as W, HV, T, J, K, and I highlight mixtures with West Eurasia, whereas F, Z, and N reflect East Eurasian connections. Remarkably, we also detected haplogroup X (0.6%), a rare lineage with a wide but patchy distribution across Eurasia. Such diversity suggests that the maternal gene pool of the Thar has been shaped by multiple sources from different regions and times (Table S17). On the paternal side, Y-chromosomal analysis also demonstrates heterogeneity. Among the Thar dwellers, the most common haplogroups were R1a (22.1%), associated with Indo-European expansions, and L1a (17.5%), which is associated with Dravidian lineages. H1a (11.2%) and R2a (13.9%) further underscores the presence of the indigenous population. At the same time, lineages such as J2a, J2b, and G2a point to introgression from Neolithic lineage and later movements from the Middle East and Central Asia, while C1b and Q1a trace connections extending into Central Asia and Siberia (Table S18).

## Discussion

Our study shows distinct genetic patterns among present-day craft artisan communities. The crafts of these communities in the northwest region are not just artistic expressions but also integral to their cultural identity and livelihoods, a legacy passed down through generations. We analyzed the broader genomic relationships and ancestry of these craft artisan communities and pastoral groups in the Thar Desert. We also analyzed their linkages with neighboring South Asian populations and global groups. Importantly, we did not classify the assigned populations based on oral history alone, but rather by their traditional occupations through endogamy and customary practices across generations.

Communities and artisan groups in the Thar illustrate historical connections through both craft and lineage that are also recognized by the Government of India (https://handicrafts.nic.in/). For instance, TH-IJE-1-MU, the gold-embossing artisans of Bikaner, trace their ancestry to Persian craftsmen invited during the Mughal era, who introduced naqashi manoti techniques and carried a legacy of Central Asian artistry. This Central Asian thread is also indicated for the pastoralist group TH-IJE-1-PR, whose nomadic lifestyle recalls origins in Baluchistan and Afghanistan, later adapting to the Thar’s ecology through long-distance seasonal mobility with caravans. The wooden craft group (TH-IJE-4-R), associated with the Rathakar, or chariot builders, doors/kawads. Their guild-based traditions in carving, measurement, and geometry echo Vedic building practices and are even mentioned in the ancient scriptures (*Skandapurana*). Similarly, the potters (TH-IJE-2-R) sustained lime kilns and pottery traditions engaged in antiquity. Their craft connects to the Namazga and Shahr-i Sokhta cultures and continues in the utilitarian pottery of the Indus Valley, also referenced in the *Yajurveda*. Collectively, these groups exemplify artisanship central to both everyday life and ritual across South Asian history. Other Thar communities preserved equally distinct traditions. TH-IJE-3-R combines agriculture, cattle-herding, embroidery, and weaving, with some also engaged in skinning and tanning. TH-IJE-1-U, descended from shoemakers, specialized in leatherwork, producing swords, saddles, belts, footwear, and other goods for warrior clans. The TH-IJE-1-SN, traditionally snake charmers and performers known as Gypsies of Rajasthan’s Deserts, lived a nomadic life and were sustained by trading snake venom, music, and dance, a heritage later recognized by UNESCO. Finally, TH-IJE-2-U are cloth printers and dyers who are renowned for their tie-and-dye textiles. They adopted this profession during the Mughal period, with some members later undergoing religious conversions.^2,8,81–85^ The settlement of these communities in the Thar Desert can be understood in relation to both environment and livelihood. While the region is harsh and resources are limited, its dry climate favored crafts that required low humidity, such as tie-dye textiles and leatherwork. Seasonal rainfall, pastoral migration, and animal husbandry provided an additional base for survival and exchange. Their craft traditions were maintained over centuries, with some techniques showing similarities to those known from the Indus Valley Civilization. At the same time, designs that bear influence from Eurasia and Central Asia reflect episodes of trans-regional movement. Royal patronage and local demand further supported the continuation of these practices. Together, these factors helped preserve cultural identity, sustain livelihoods, and tie the communities both to their desert setting and to wider networks of movement and interaction.

Present-day northwest India was historically influenced by successive waves of migration and incursions, including those of the Greeks, Scythians, Parthians, Kushans, and Huns. Taken together, these communities embody the Thar as a corridor of migration, cultural exchange, and ecological adaptation. Their hereditary crafts, anchored in oral traditions, Vedic references, Mughal patronage, and transregional linkages, remain inseparable from their genetic history.^86–90^ Previous studies on populations from southern and central Rajasthan, including both tribal and caste groups, have reported higher average heterogeneity values based on autosomal STR analyses. These groups showed stronger affinities with Indian populations, particularly North Indians, rather than with populations from the western frontier. Some tribal groups from the northwest were also found to share greater genetic similarity with Dravidian populations. In addition, populations such as Gujjar and Kambhoj, located near ancient Rajasthan sites of civilization, have been studied for their affinities with both ancient and modern populations. These populations are also included in our study for comparison and reference^.36,68,91–94^

From multiple genetic distance analyses, we found that craft artisan communities aligned with the Indo-European cline of South Asia, and there was substantial heterogeneity within the northwest region, which maintained separate and distinct population groups over a long period. Genetic differentiation was found to be lower in the population from the Gangetic Plain and the western frontier, that is, proximal to the Indus. We found that the Thar artisan population shows affinity with Sindhis; however, the populations like Pastoralists and artisans (Woodcarvers and Persian gold embossers) of the Thar artisans share more affinity with western frontier populations like Pathan, which shows a Central Asian connection. This is reflective of the close geographical proximity of Sindh to the Thar Desert, wherein Sindhis were merchants and traders of goods and crafts. This suggests that population structure experienced mobility along regional routes during the migration of Indo-Europeans toward the Gangetic Plain after the decline of civilization. These artisan and pastoralist groups show a higher degree of West Eurasian genetic influence, comparable to or even exceeding that observed in other North Indian populations. This pattern is consistent with previous studies which indicate that northwestern Indian and upper-caste populations across India share greater genetic similarity with groups from Central and West Asia, as well as parts of Europe.^21,36,68,77,95–97^ In contrast, within the Thar region, artisan communities such as performing artists, potters, and textile and leather artists showed stronger genetic affinity with local Indian populations and diverse South Asian ethnic groups, while showing relatively lower affinity with West Eurasian populations.

Our observation is further supported by the proportion of ANI (Ancestral North Indian), now recognized as ANE (Ancestral North Eurasian),^98^ which was consistently higher in most of the Thar artisan groups, comparable to levels observed in Kalash and Pathan populations, while being lowest in TH-IJE-1-SN. Given that ANI ancestry is widely shared with Central Asians, Caucasians, Middle Easterners, and parts of Europe, these results underscore substantial variation in West Eurasian genetic influence among artisan communities of the Thar region. These ties to Steppe pastoralism and Indo-European expansions in India are also reflected in the dominance of R1a of Y-chromosome haplogroups among many craft artisans. These are in addition to the other Middle Eastern and Central Asian associated haplogroups, such as J2a, J2b, and G2a. Noteworthy, the presence of the Y chromosome L1a suggests connections to Neolithic migrations, possibly linked to the spread of Dravidian languages and agriculture from Iran into South Asia, corroborating the Elamo-Dravidian hypothesis. Also, the highest frequencies of South Asian-related haplogroup M amongst the mtDNA haplogroups indicate a strong local or regional maternal lineage. Noteworthy, in the three Thar populations prominently in TH-IJE-2-U, with links to West Eurasia, we also observe the consistent presence of U7, W, and R2, which further substantiates our observations from autosomal studies.

The observed pattern of heterogeneous population structure and genetic drift in several Thar populations, particularly migrant artists and tie-dye artists, reflects broader founder events documented across South Asia. These patterns may be linked to historical episodes of migration and religious conversion occurring within the last 400–500 years. The observed homozygosity patterns, corroborated by runs of homozygosity (ROH) and effective population size estimates, indicate isolation and genetic drift due to endogamy and geographical constraints. These demographic patterns show consolidation of artisan communities under guild-based production systems that became increasingly rigid during the Mughal period, when craft specialization was institutionalized within hierarchical social structures.

Earlier genetic studies have shown that the populations of the Indian subcontinent fall along a broad cline shaped by two major ancestral components. Ancestral South Indians (ASI) derive from indigenous South Asian hunter-gatherers and ancient Iranian Neolithic farmers, while Ancestral North Indians (ANI) reflect ancestry from Middle to Late Bronze Age (MLBA) steppe pastoralists connected to Eurasia. Situated along a historical corridor in northwestern India, the Thar Desert populations show varying proportions of MLBA steppe ancestry, with some northwestern groups (NWI) carrying the highest levels reported in India to date.^20–22^ These findings are supported by multiple lines of evidence, including D-statistics, outgroup f3-statistics, and admixture modeling with qpAdm.

The presence of these ancestral components likely reflects both proximal and distal migration events. Elevated allele sharing in the Northwest region highlights the Thar’s role as a genetic crossroads, shaped by gene flow from diverse external sources. Their ancestry shows connections to the Iranian plateau and interactions with steppe groups, Indus Valley populations, and later diasporas. In addition, it further includes Mesolithic hunter-gatherer ancestry (EEHG, CHG), Neolithic farmer contributions from Anatolia and Iran-Turan, and later Bronze, Copper, and Iron Age influences from the Central Asian steppe and adjacent regions.^99,100^ These genetic inputs, particularly in Thar, align with the spread of traits like lactase persistence (−13910*T allele), introduced via pastoralist migrations from the Middle East or Anatolia around 5,000–3,000 years ago.^69,70^ The geographic distribution of lactase persistence across India exhibits a clear northwest-to-southeast declining pattern and heterogeneity in its frequency among populations of the Thar Desert. We found that Ror and TH-IJE-1PR show the strongest signals in South Asia, likely linked to their traditions of animal husbandry, agro-pastoralism, and dairy consumption. These groups share ancestry from the Anatolia Neolithic (Anatolia N) and Caucasus Hunter-Gatherers (CHG).^43^ In contrast, another pastoralist group from the same lineage, IENLPX4, shows a stronger affinity with Indian-specific ancestry, while the pastoralist group in our study displays greater West Eurasian-related affinity. Specifically, it leans more toward Neolithic Iranian and BMAC lineages (∼20 to 25% Anatolia_N-related ancestry), which are geographically and temporally closer to the Indus Valley populations and the Indo-Iranian frontier. A complementary signal of West Eurasian influx is also reflected in the frequency of the SLC24A5 rs1426654 varian**t** (associated with lighter skin pigmentation).^72,73^ This allele occurs at higher levels in most of the Thar population and in some Northwest Indo-European-speaking groups, but its frequency decreases substantially toward the south.

In conclusion, our study positions the Thar Desert at a remarkable crossroads where geography, migration, craft traditions, cultural practices, and ecology converge to shape the genomes of its distinct craft artisan communities. Beyond social isolation, endogamy, and consanguinity, the artistic traditions integral to cultural identity and livelihood also contribute to genetic distinctiveness. The genomes of the artisan and pastoralist groups thus act as a palimpsest, preserving signals from successive demographic events. Some communities show higher genetic drift and population isolation, with the timing of founder events aligning with episodes of social and political change across the medieval, early modern, and colonial periods. Religious practices, occupational shifts driven by need, community separation, and the formation of new social groups through occupational mobility all contributed to their genetic distinctiveness. Endogamy, maintained for socioeconomic and cultural reasons, reinforced group boundaries and shaped their genetic structure in recent centuries. When integrated into broader Indian and global genomic datasets, these populations trace their roots both to movements along the western corridor and to settlements from the Indian interior, reflected in clinal genetic patterns. They preserve layered imprints of demographic episodes, from early Neolithic farming expansions to Bronze Age steppe-related movements and later exchanges. Thar populations also show examples of a gene–culture co-evolution differentiated by selective pressures that leave genetic signatures on adaptive traits such as lactase persistence and skin pigmentation. The diversity of uniparental markers, spanning South Asian lineages alongside inputs from West and Central Eurasia, further underscores the heterogeneity and fine-scale structure of the Thar. Although the current dataset provides valuable insights about Thar, the inclusion of higher-density genomic data and the discovery of ancient samples from this region would enable a more refined reconstruction of Thar’s genetic history. This study thus provides the first genomic baseline for the Thar Desert populations. Beyond reconstructing population history, this baseline will be valuable for detecting convergent selective signatures in extreme environments, assessing the burden of rare diseases arising from founder events, and for complex disease risk stratification, particularly pertaining to living in the desert.

## Acknowledgements

We gratefully acknowledge the support and cooperation of artisans during this study, as well as the funding provided by the Jodhpur City Knowledge and Innovation Foundation (JCKIF) and a seed grant to MM and the HPC facility at IIT Jodhpur (JCKIF/Thar/Proj-01/2022, I/SEED/MTM/20220130). Genotyping support from CSIR-Institute of Genomics and Integrative Biology (CSIR-IGIB) is also acknowledged. We are particularly thankful to Dr. Pooja Sharma for their valuable technical support. We also express our gratitude to Dr. Arindam Mitra, National Institute of Biomedical Genomics (NIBMG), Kolkata, for assistance with preliminary genotype data processing and curation. We extend our thanks to the Department of Bioscience and Bioengineering, IIT Jodhpur, and to the students for their participation. We acknowledge the contributions of Dr. Manasi Mukherjee (Project Coordinator) and Dr. Ravi Pratap Singh for participating in the health camp and field survey. Thanks to Mr. Padam Singh and Mr. Jitendra Singh for their work as field assistants. Finally, we acknowledge the financial assistance received in the form of student fellowships from the University Grants Commission (UGC) to RJ and the Ministry of Human Resource Development (MHRD), India, to SMR.

## Author’s contributions

MM conceived the study. GNJ helped in the population identification and community engagement. RJ and SMR performed the experimental work and genotyping. MF helped in the genotyping experiments. RJ, SMR, DJ, MM, and GNJ conducted field visits and carried out the anthropological survey and engagement, health camp and data collection. RJ performed the population data analysis with input from PJ and MM, and SMR also contributed to the analyses. RJ, SMR, and MM wrote the manuscript. All authors reviewed and approved the final version.

## Web resources

The 1000 Genomes Project, http://www.internationalgenome.org/home

HapMap3, https://www.sanger.ac.uk/resources/downloads/human/hapmap3.html

PhyloTree, http://www.phylotree.org/

HGDP Sequence: https://www.internationalgenome.org/data-portal/data-collection/hgdp

AADR v54, https://dataverse.harvard.edu/dataset.xhtml?persistentId=doi:10.7910/DVN/FFIDCW

## Data and code availability

The datasets and code supporting this study have not been deposited in a public repository because they contain sensitive population-genetic data and participant information that could compromise individual privacy. However, they are available from the corresponding author upon reasonable request.

## Supplementary Figures

**Figure S1:**
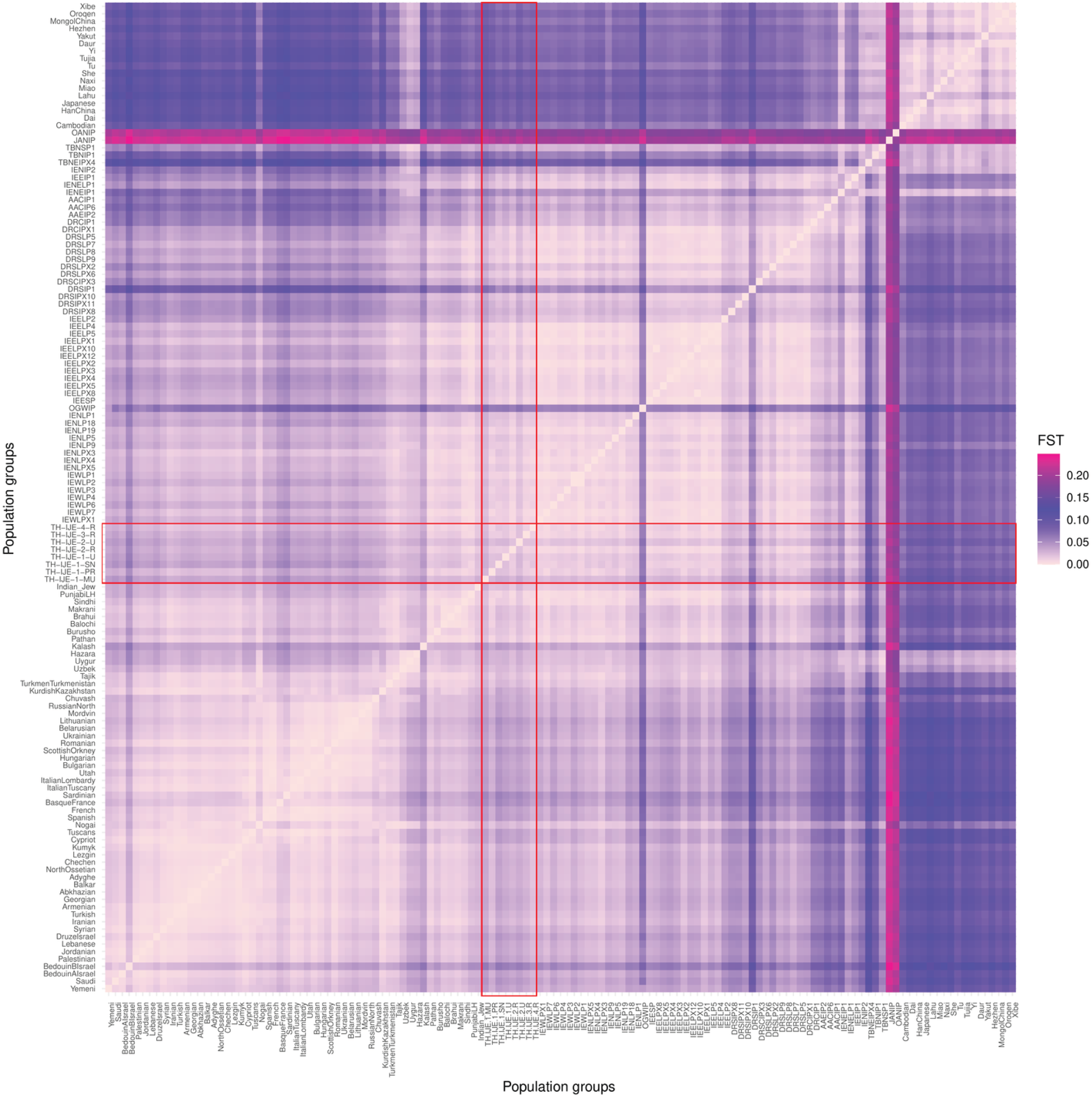
Pairwise population differentiation measured by Fst. Matrix of pairwise Fst (fixation index) values calculated using SMARTPCA across all reference populations. Shading intensity reflects the magnitude of genetic divergence, as indicated by the color scale. Populations are ordered geographically and grouped by continent, with the newly sampled Thar Desert craft populations highlighted for comparison.

**Figure S2:**
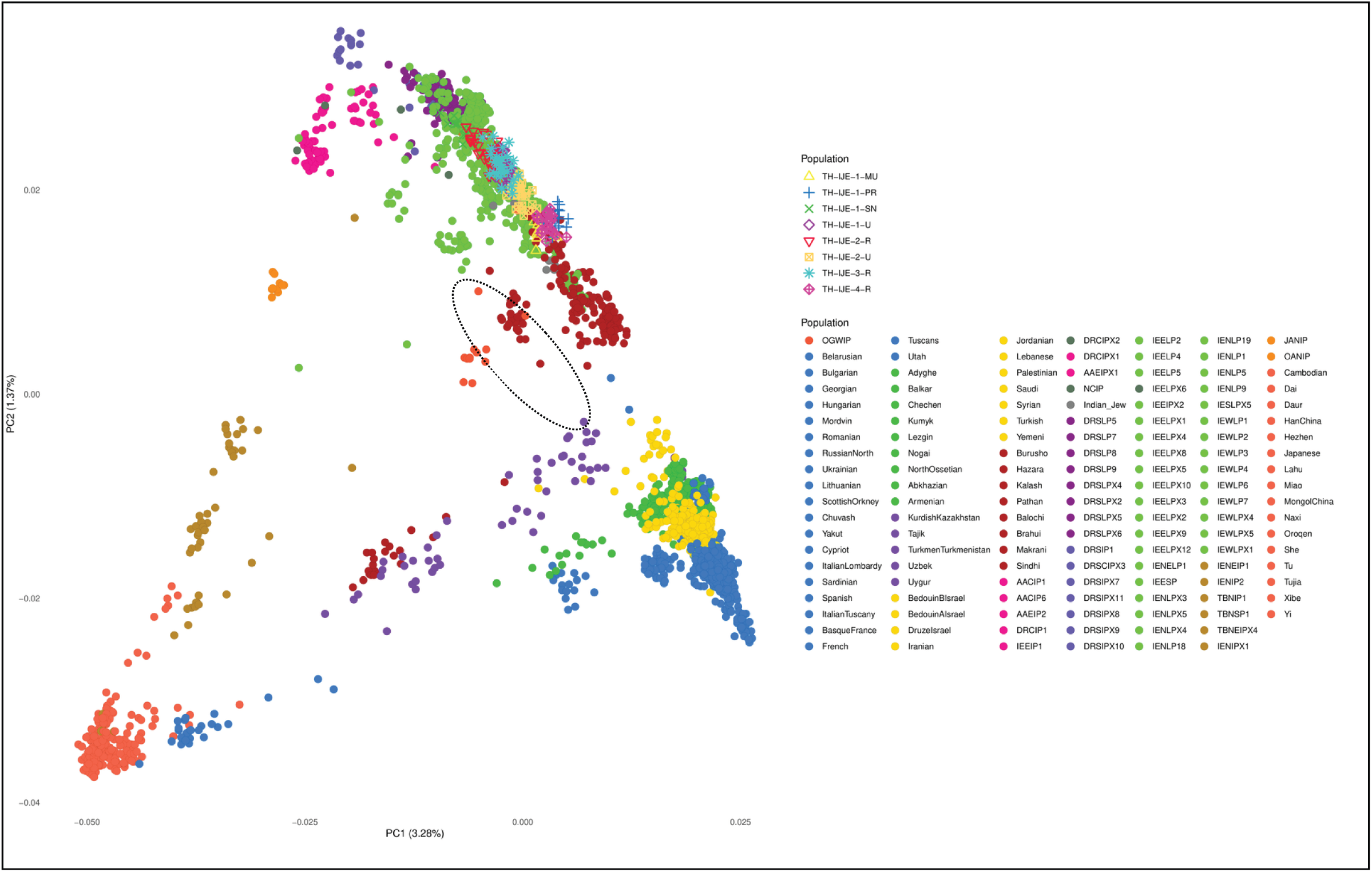
Genome-wide principal component analysis. Genome-wide principal component analysis (PCA) showing the genetic structure of Thar Desert craft populations in the context of modern South Asian and other Eurasian populations. Populations are ordered by geography and linguistic affiliation and are color-coded accordingly. The PCA highlights the distribution of study populations along the Ancestral North Indian (ANI)–Ancestral South Indian (ASI) cline, illustrating their relative positions within South Asian genetic diversity.

**Figure S3:**
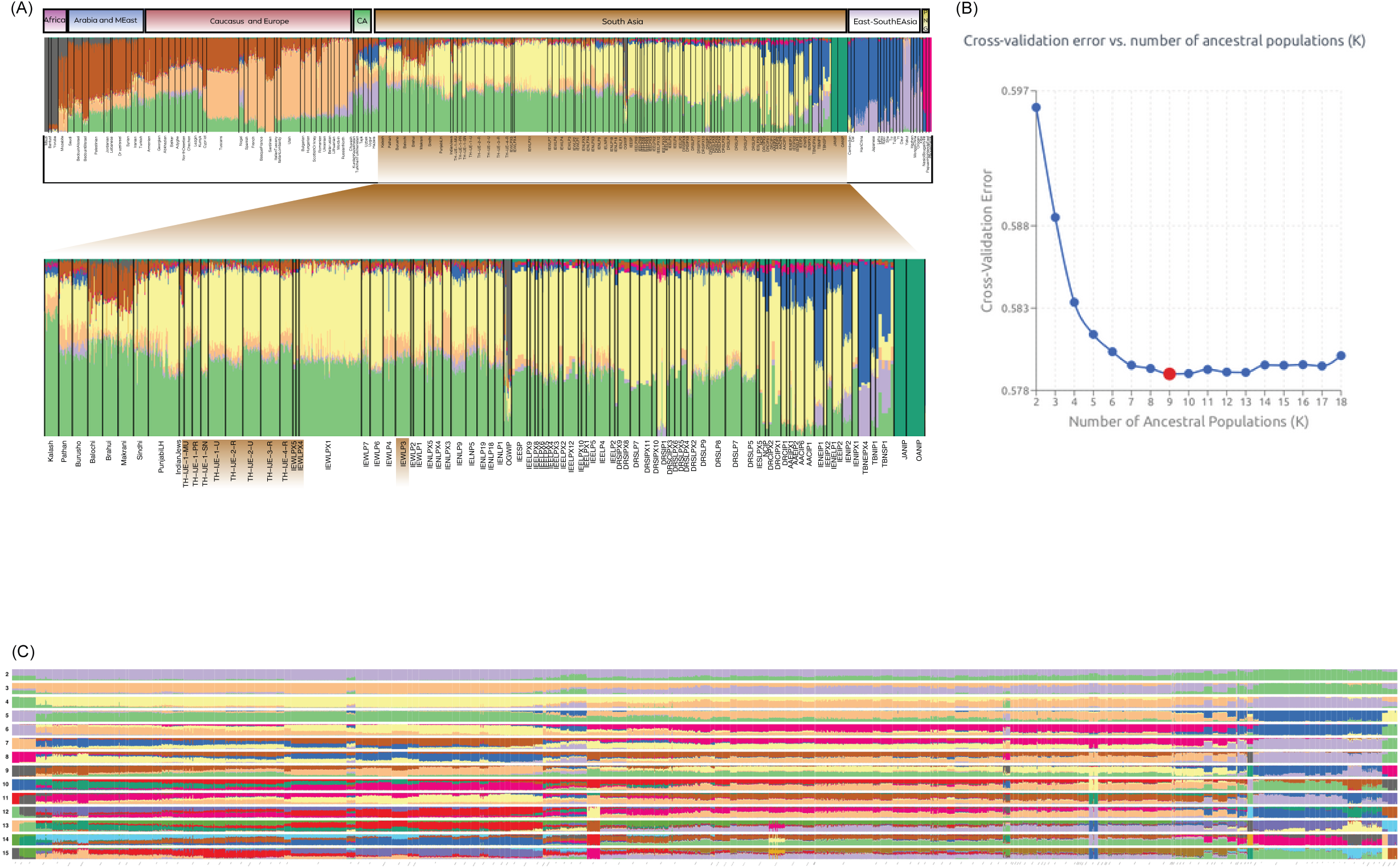
Genetic ancestry components inferred from ADMIXTURE analysis. (A) Unsupervised clustering results at K = 9, displayed as a stacked bar plot. Global populations are ordered geographically (Africa, Arabia and the Middle East, Central Asia, Caucasus and Europe, Southeast and East Asia, Papua New Guinea(PNG), and South Asia). The inset highlights South Asian populations, with the newly studied Thar Desert groups labelled (see Methods).(B) Cross-validation results for ADMIXTURE runs with K values ranging from 2 to 18, illustrating the hierarchical structure of genetic components across populations.(C) ADMIXTURE results across multiple K values (K = 2–15), shown as stacked bar plots, illustrating the hierarchical emergence of ancestry components.

**Figure S4:**
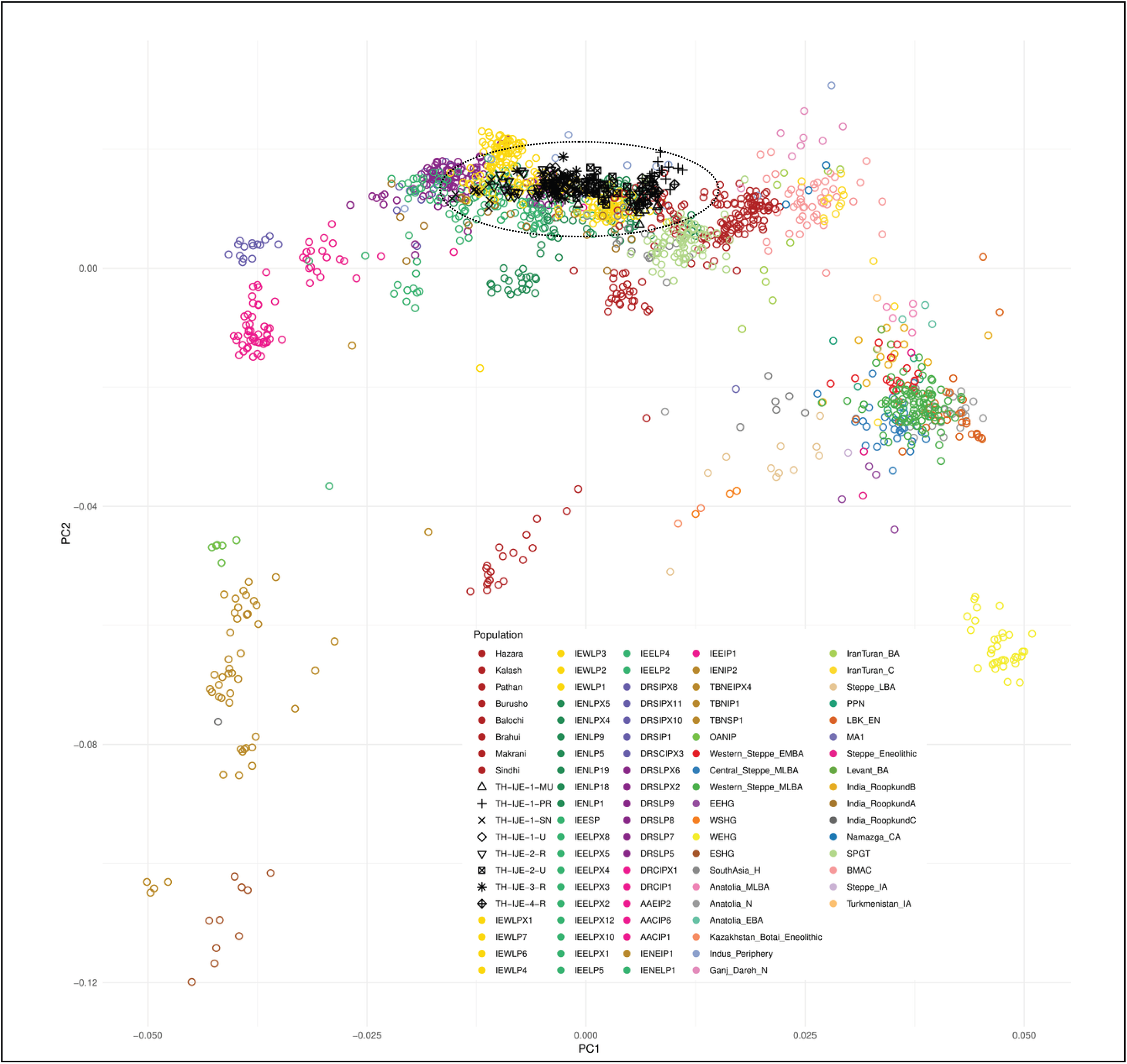
Genome-wide principal component analysis. Genome-wide principal component analysis (PCA) illustrating the genetic structure of Thar Desert craft populations in the context of ancient DNA. The plot includes modern Eurasian populations alongside selected ancient sources from the Central and Western Steppe, Iran, Anatolia, Central Asia, and South Asia, highlighting the study populations in relation to both ancient and contemporary South Asian

**Figure S5:**
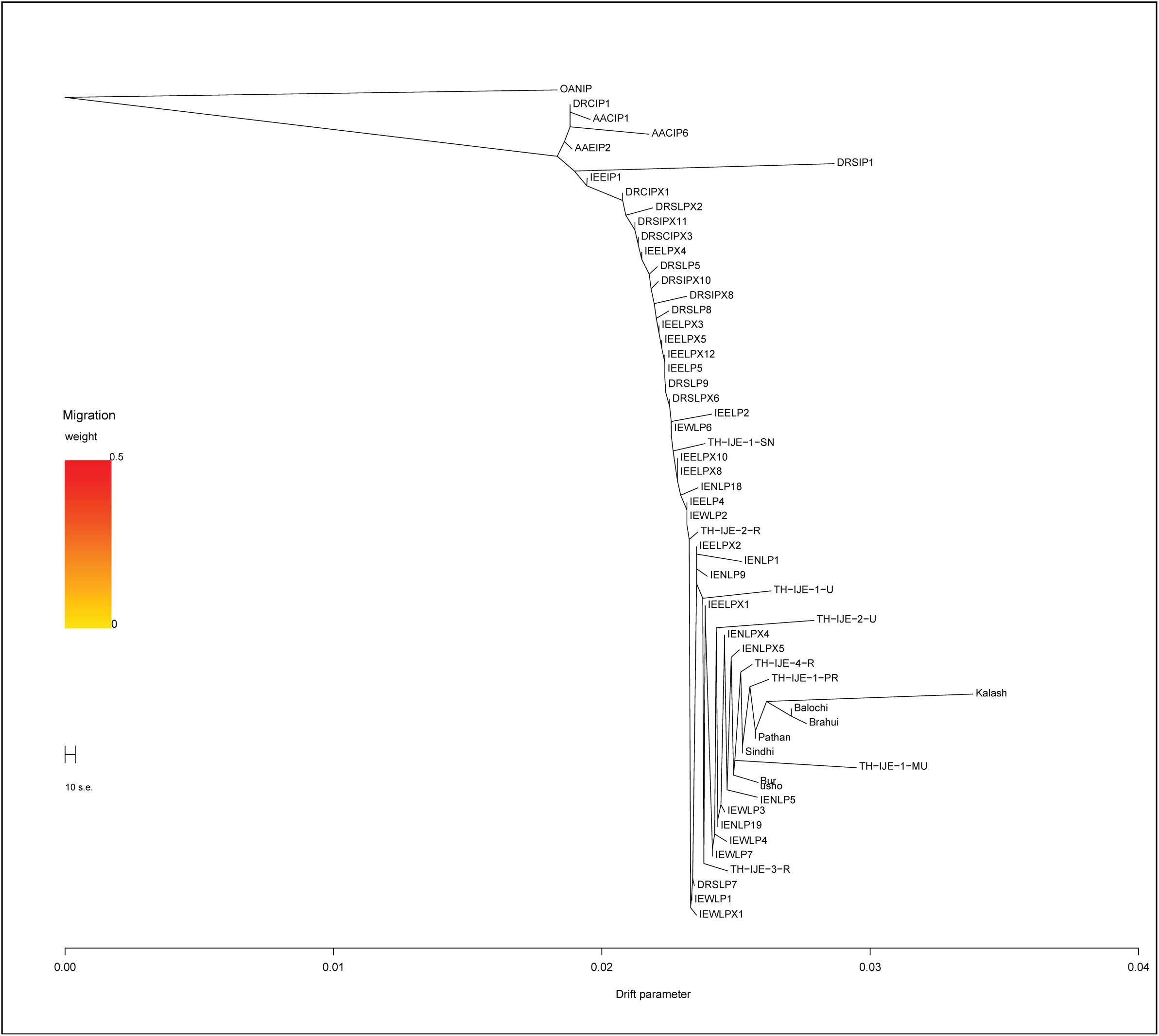
Treemix maximum likelihood tree. Maximum likelihood tree depicting the genetic relationships of Thar Desert populations with neighboring groups, including the Western Frontier and other Eurasian populations, as inferred by TreeMix. Branch lengths reflect population-specific drift, highlighting higher genetic drift observed in Thar populations relative to other groups.

**Figure S6:**
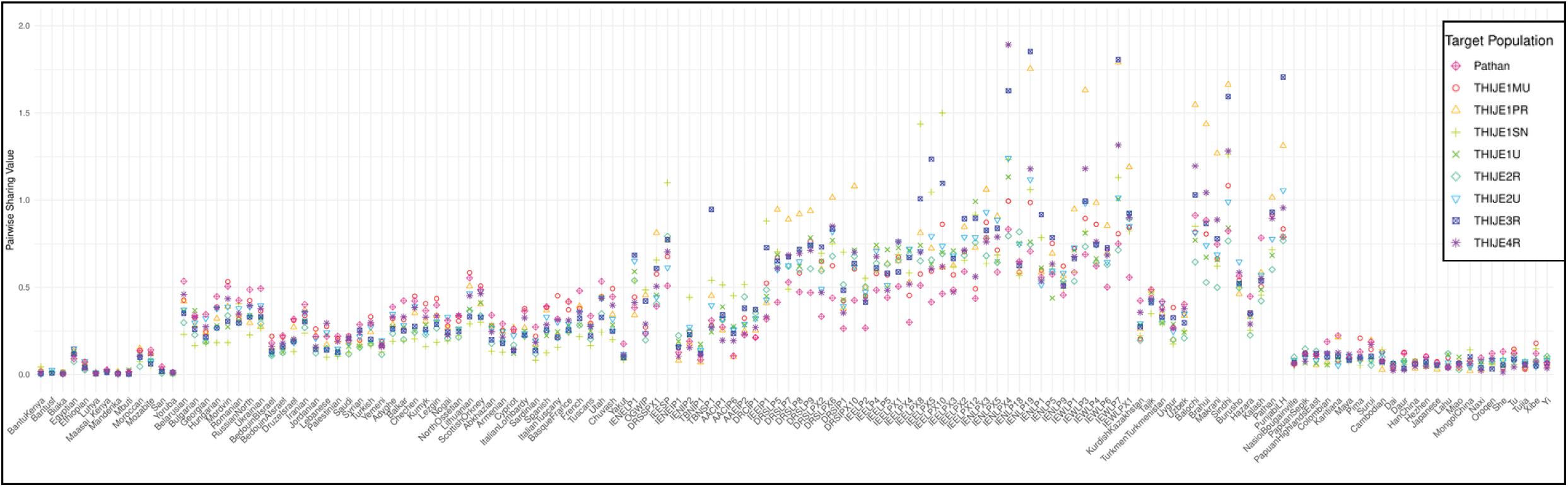
Identity By Descent sharing in Thar and other population. Average number of IBD (Identity-By-Descent) segments shared per pair of individuals in Thar Desert populations and the Pathan ethnic group from the Western Frontier, as inferred using refined IBD (see Methods).

**Figure S7:**
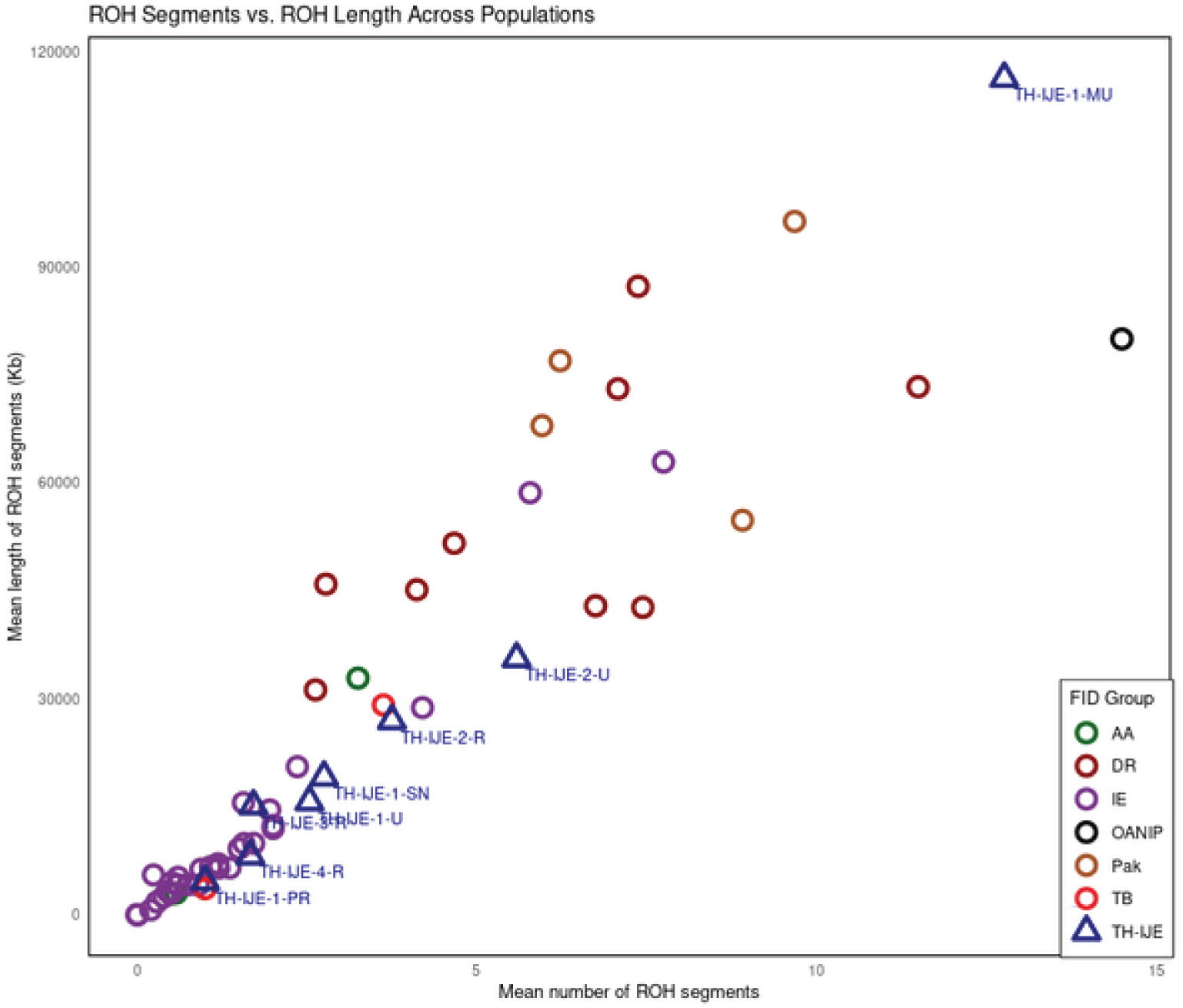
Runs of Homozygosity in Thar and other South Indian populations. Runs of Homozygosity (RoH) in Thar Desert populations compared with other South Asian groups. The plot displays the mean number of RoH segments versus the mean segment length in kb. RoH were identified using PLINK v1.9 (see Methods). Populations are colour-coded and grouped by linguistic affiliation. The distribution patterns of recent inbreeding and population isolation in Thar populations relative to other South Asian groups.

**Figure S8:**
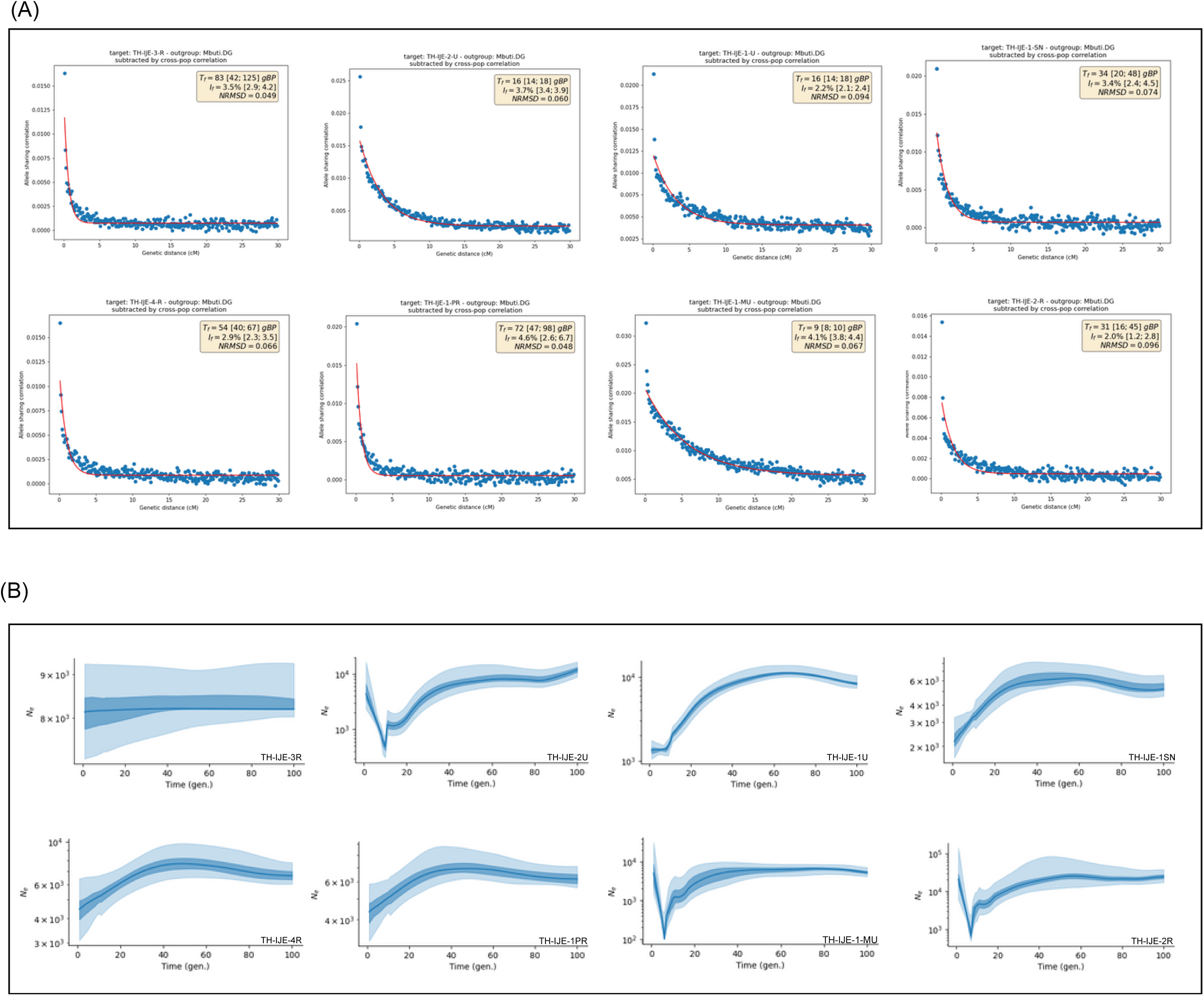
Demographic history of Thar Desert populations through Founder events. (A) Founder events inferred using ASCEND, showing the occurrence and magnitude in the Thar population. Plots depict the time of founder events (Tf) and their strength (If), highlighting any historical bottlenecks or reductions in population size.(B) Historical effective population size (Ne) inferred using HapNe-IBD. The x-axis represents generations before present, and the y-axis shows Ne estimates. Light and dark shaded areas correspond to 95% and 50% confidence intervals, respectively, obtained via bootstrap quartiles, capturing recent changes in the effective population size of the Thar populations.

**Figure S9:**
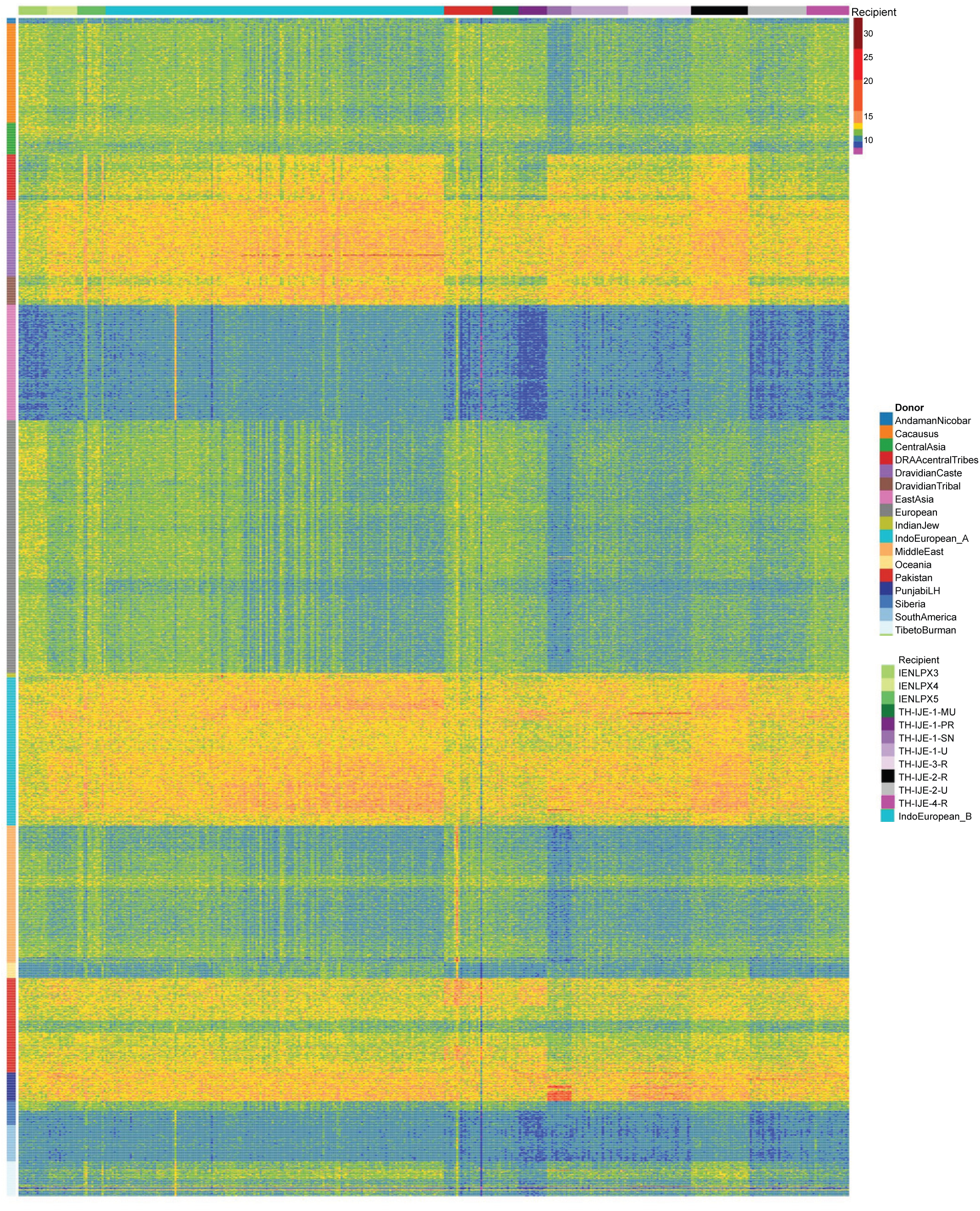
Heatmap of mean haplotype chunks in Thar. Heatmap showing the mean haplotype chunks received by Thar Desert populations and selected North/Western Indo-EuropeanA populations, as inferred using CHROMOPAINTER. Chunks represent shared genomic segments donated from populations across Eurasia, highlighting patterns of allele sharing and gene flow, particularly into the Thar populations. The color scale reflects the relative amount of shared ancestry, illustrating contributions from different Eurasian source populations to both the study and comparative groups.

**Figure S10:**
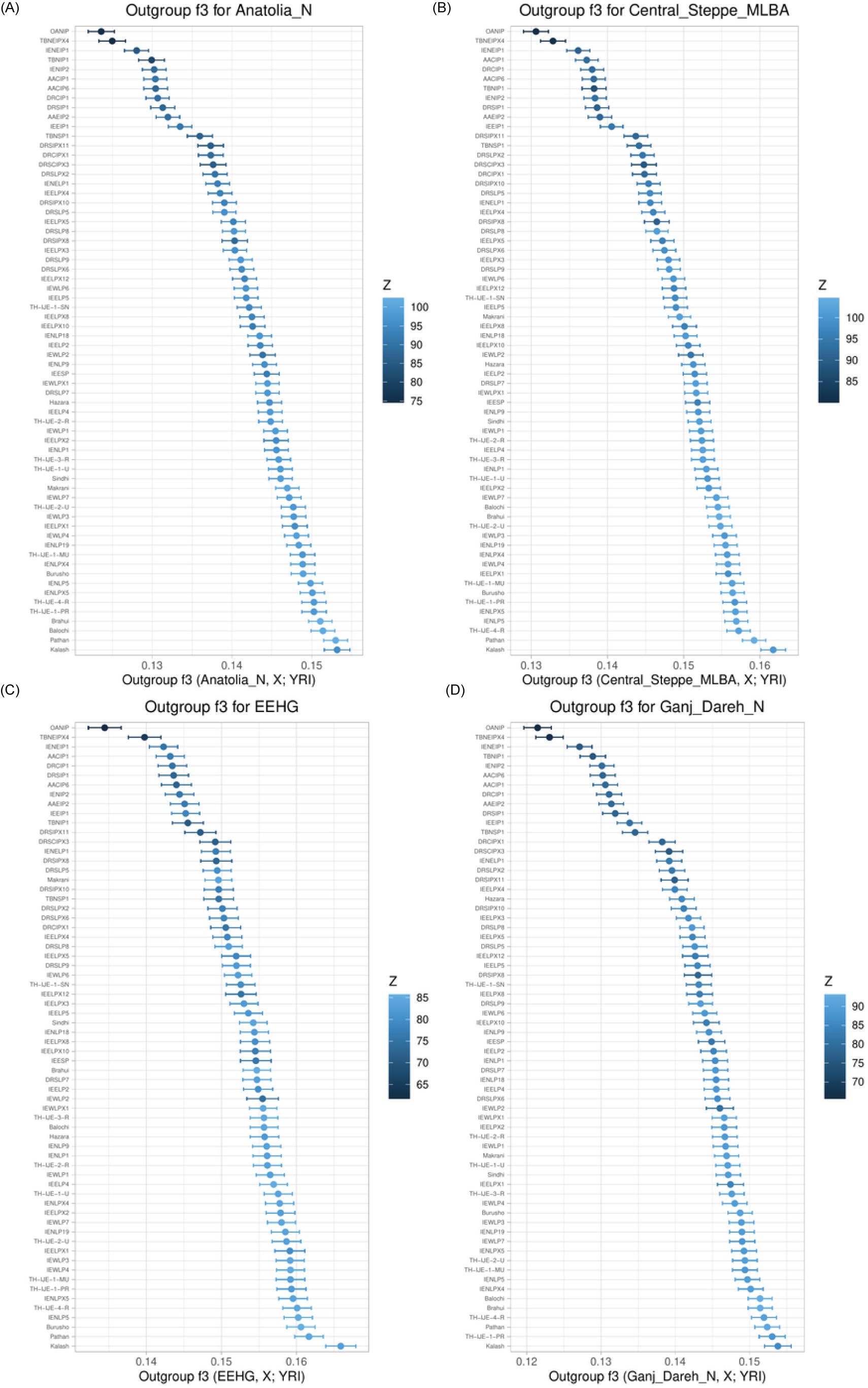
f₃ statistics of Thar and South Asian population with the ancient population. Outgroup f₃ statistics measuring shared genetic drift between populations relative to an outgroup, using ancient reference sources for South Asian and West Eurasian populations. Panels show results with (A) Anatolia_N, (B) EEHG, (C) Central_Steppe_MLBA, and (D) Ganj_Dareh_N. The statistic is calculated as f₃(Ancient Source, X; Yoruba), where X represents a South Asian or West Eurasian population. Error bars indicate jackknife standard errors. Colors correspond to Z-scores, with the gradient legend shown alongside.

